# Variation of C-terminal domain governs RNA polymerase II genomic locations and alternative splicing in eukaryotic transcription

**DOI:** 10.1101/2024.01.01.573828

**Authors:** Qian Zhang, Wantae Kim, Svetlana Panina, Joshua E. Mayfield, Bede Portz, Y. Jessie Zhang

**Affiliations:** Department of Molecular Biosciences, Austin, Texas, 78712; McKetta Department of Chemical Engineering, University of Texas, Austin, Texas, 78712; Department of Pharmacology, Chemistry, and Biochemistry, and Cellular and Molecular Medicine, University of California San Diego, La Jolla, California 92093; Dewpoint Therapeutics, 451 D Street, Boston, Massachusetts 02210

**Author notes:** Contact should be addressed to Y. Jessie Zhang.

**Keywords:** Transcription regulation, post-translational modification, RNA polymerase II, Liquid-liquid phase separation, and alternative splicing

## Abstract

The C-terminal domain of RPB1 (CTD) orchestrates transcription by recruiting regulators to RNA Pol II upon phosphorylation. Recent insights highlight the pivotal role of CTD in driving condensate formation on gene loci. Yet, the molecular mechanism behind how CTD-mediated recruitment of transcriptional regulators influences condensates formation remains unclear. Our study unveils that phosphorylation reversibly dissolves phase separation induced by the unphosphorylated CTD. Phosphorylated CTD, upon specific association with transcription regulatory proteins, forms distinct condensates from unphosphorylated CTD. Function studies demonstrate CTD variants with diverse condensation properties in vitro exhibit difference in promoter binding and mRNA co-processing in cells. Notably, varying CTD lengths lead to alternative splicing outcomes impacting cellular growth, linking the evolution of CTD variation/length with the complexity of splicing from yeast to human. These findings provide compelling evidence for a model wherein post-translational modification enables the transition of functionally specialized condensates, highlighting a co-evolution link between CTD condensation and splicing.

## Introduction

The C-terminal domain of the largest subunit of RNA polymerase II (CTD) is a highly disordered region found in RNA polymerase II (Pol II), the workhorse responsible for transcribing all protein-coding mRNAs as well as some small nuclear and microRNAs in eukaryotes^1,2^. Different transcriptional regulatory proteins are recruited to RNA Pol II via the CTD to facilitate the progression of transcription^3^. The CTD recruits transcriptional proteins through extensive post-translational modifications (PTMs), with phosphorylation being the key modification during active transcription^4,5^.

The sequence of the CTD is surprisingly simple, with consensus heptads (historically numbered as Y_1_S_2_P_3_T_4_S_5_P_6_S_7_) repeated many times dependent on the species (e.g., 26 in *S. cerevisiae* and 52 in humans)^1^. Despite of the simplicity, five of the seven residues in the heptad repeats are subject to phosphorylation, and the two proline residues can undergo isomerization, which can affect recognition by different CTD-interacting domains (CIDs) of diverse proteins that dynamically associate with Pol II throughout the transcription cycle^6,7^. Therefore, this simple repetitive sequence possesses an enormous capacity to encode information via combinatorial phosphorylation^8^. Different residues of the heptad repeats on CTD get phosphorylated at various stages of transcription, with Ser5 as the major species during initiation, recruiting capping enzymes^9^, and Ser2 at elongation/termination recruiting splicing and termination factors^1^. Instead of the traditional view of the CTD as scaffold for protein binding, the paradigm has shifted in recent years towards an ensemble view whereby Pol II functions within transcriptional condensates, the composition of which is governed in part by CTD phosphorylation^10–13^.

Mounting evidence shows that the CTD drives the Pol II participation into condensates^10–12,14,15^. At transcription initiation, RNA polymerase II accumulates in LLPS droplets with components of the Mediator like MED1^10–12,15^. A report observes RNA polymerase II also enters droplets characterized by spliceosome components in the same gene loci^11^. Mounting evidence suggests Pol II condensation via the CTD is not simply phenomenological, but evolutionarily adaptive. For example, aberrant CTD condensation properties lead to developmental failures in Drosophila, and cold tolerant fungi tune CTD condensation and its regulation for environmental adaptation^10,16^.

Post-translational modifications (PTMs), such as phosphorylation, arise as an important mechanism trigger that alters condensate states. Diverse outcomes have been reported upon protein phosphorylation on IDRs, which are frequent targets for PTMs. Both dissolution and nucleation of condensates has been demonstrated in response to PTMs^17–19^. Phosphorylation on the CTD is particularly interesting since its different phosphorylation states govern the recruitment of transcriptional regulators^2,3,20^. Stereospecific binding of regulators, and emerging view of the CTD as a modulator of transcriptional condensate formation, suggest a model of transcriptional condensates sorting and enriching CTD interacting factors that in turn bind preferred heptad motifs with preferred patterns of phosphorylation. We thus focus on interplay between CTD phosphorylation, condensate formation, and CTD-interacting protein recruitment and their effects on transcription.

We employed a suite of CTD mutants, a series of CTDs with distinct post-translational modifications (PTMs) applied enzymatically, and a set of CTD interacting proteins recognizing discrete patterns of CTD PTMs to derive rules of PTM and protein binding that govern CTD condensate formation and topological organization. Our findings reveal that CTD phosphorylation reversibly dissolves CTD condensates, yet binding by CTD interacting partners antagonizes phosphorylation mediated condensate dissolution. We demonstrate that phospho-CTD condensates are enabled by the binding of phospho-specific CTD interacting proteins, and that factor binding can generate CTD condensates with layered topologies characterized by an unmodified CTD core and phospho-CTD shell harboring CTD interacting partners. The transcriptomic analyses of human cells expressing RNA Pol II mutants harboring CTDs with varied abilities to form condensates reveal differences in promoter binding, alternative splicing, and ultimately growth defects that corroborate rules of CTD condensation derived from our biochemical assays. Our work establishes a biochemical principle for the maturation of RNA Pol II condensates associated with distinct transcriptional stages and support a model of RNA Pol II flux from promoter-associated initiation condensates to elongation condensates that support the co-transcriptional splicing. In addition, we also describe an intrinsic and essential function of the variation of CTD in regulation of cell survival through altering exons skipping and intron retaining of alternative splicing events.

## Results

### Reversible phosphorylation of the CTD leads to phase transition

We employed a well-established system to investigate LLPS of RNA polymerase II in vitro. This involved purifying a GST-tagged *S. cerevisiae* CTD domain containing 26 heptad repeats, predominantly composed of the consensus sequence (yCTD, Figure S1A)^12^. To ensure the robustness and rigor of our biochemical analysis, we ensured the purity of all protein samples used in the studies, including both the CTD variants and CTD binding proteins, through gel filtration chromatography, SDS-PAGE, differential interference contrast microscopy and light scattering. To provide further evidence of sample purity, the CTD binding proteins used in the study were able to produce diffracting quality crystals.

We covalently attached a fluorophore to the N-terminus of the recombinant GST-CTD for visual detection. Confocal microscopy and phase diagram reveal that unphosphorylated CTD forms concentration-dependent condensates in the presence of a crowding agent, dextran (Figures S1B), behaving as liquid droplets with the ability to fuse, as reported previously (Figure S1C)^12,14^. We next conducted experiments to test whether phosphorylation by CTD kinases affects the ability of the CTD to form condensates^21,22^. Two kinases (Erk2 and Dyrk1a, respectively) were used to generated CTDs distinct patterns of phosphorylation of Ser5^23^ (called pSer5 CTD in the rest of the paper) and phosphorylates Ser2^24^(pSer2 CTD) in the context of the consensus sequence. Kinase specificity was confirmed with single, double, triplet, and 26-CTD repeats as substrates by Ultraviolet Photodissociation Mass Spectrometry (UVPD) ^23–25^. We conducted the kinase reactions ―treated by Erk2 or Dyrk1a ― on CTD droplets in a time-dependent manner, simultaneously monitoring the CTD phosphorylation process and condensation states (Figure 1A and 1B). The kinase phosphorylation of the CTD was monitored at each time point using Electrophoretic Mobility Shift Assay (EMSA) and Matrix Assisted Laser Desorption/Ionization – Time of Flight (MALDI-TOF) mass spectrometry (Figure 1C and Figures S1D and S1E). Simultaneously, we observed the condensate disappearance through visualization using fluorescence microscopy (Figure 1A) and turbidity assays (Figure 1B). The signal in turbidity assay decreased substantially more rapidly as an increasing amount of kinases resulted in faster phosphorylation (Figure 1B). Notably, neither kinases by themselves exhibit any condensate formation under the reaction conditions (Figure S1F). Furthermore, control conditions to exclude each phosphorylation reaction component (ATP, Mg^2+^ ion, and kinase) systematically reveal that condensate dissolves only when all kinase reaction components are present for phosphorylation to occur (Figure S1G). Rigorously, we showed that the phosphate groups placed on the CTD induce condensate dissolution.

**Figure 1.**
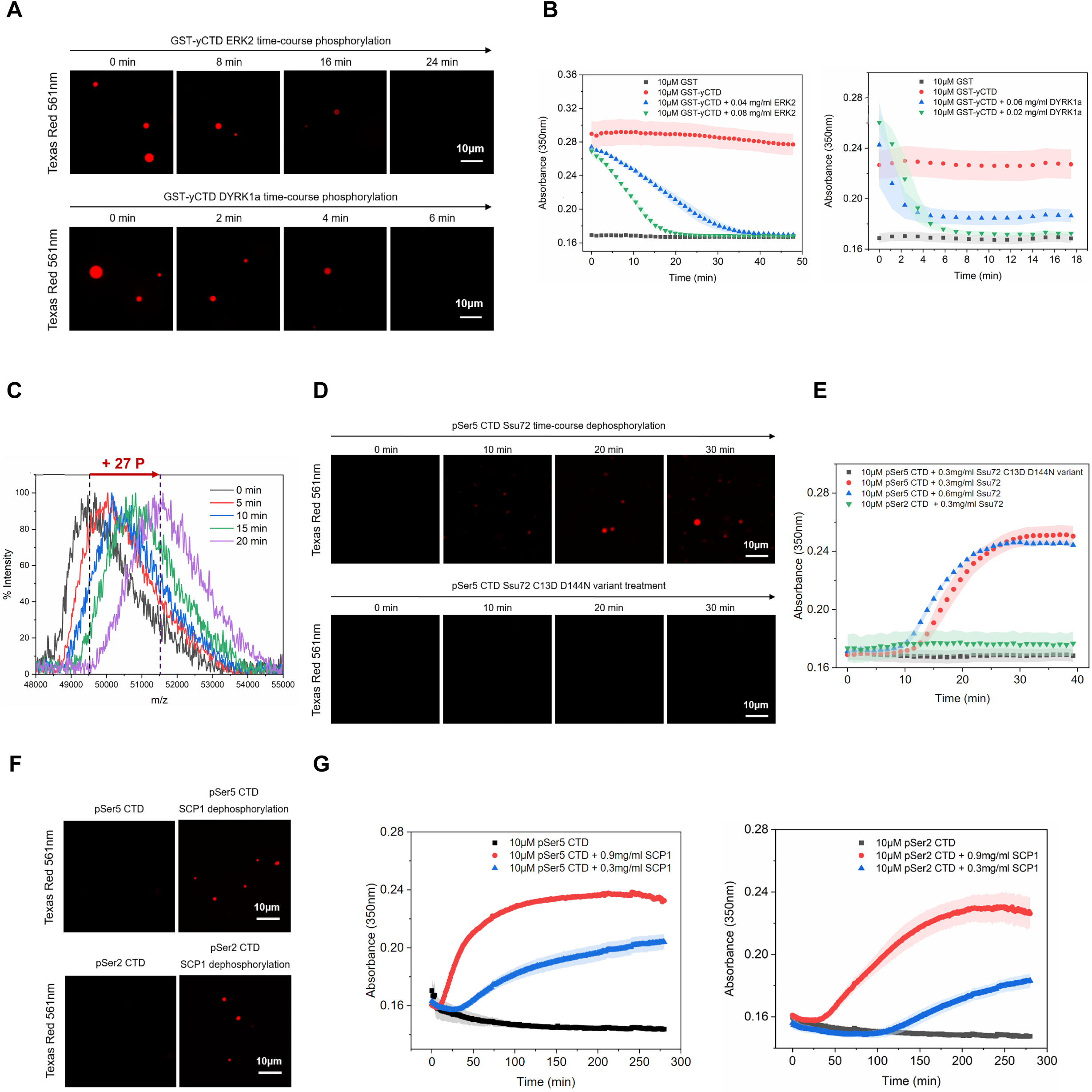
Reversible phosphorylation of the CTD leads to phase transition. (A) Phosphorylation by ERK2 and DYRK1a gradually dissolves condensates formed by unphosphorylated CTD labeled with Texas Red-X fluorophore. (B) Time-course turbidity assay of unphosphorylated GST-yCTD samples treated with ERK2 (left) and DYRK1a (right) in the presence of 16% dextran. Data points represent mean values of three replicate experiments, and error bars show the standard error. (C) MALDI-TOF MS spectrum of ERK2 treated CTD samples (pSer5 CTD) taken from turbidity assay experiments at different timepoints. (D) Time-course confocal images of pSer5 CTD (labeled with Texas Red-X) treated by Ssu72 phosphatase (top) and Ssu72 C13D/D144N variant (bottom). (E) Time-course turbidity assay of pSer5 CTD samples treated with Ssu72 in the presence of 16% dextran. Data points represent mean values of three replicate experiments, and error bars show the standard error. (F) pSer5 CTD sample (top) and pSer2 CTD sample (bottom), both labeled with Texas Red-X, are treated by SCP1 overnight prior to confocal imaging. (G) Time-course turbidity assay of phosphorylated CTD samples treated with SCP1 in the presence of 16% dextran. Data points represent mean values of three replicate experiments, and error bars show the standard error.

To interrogate whether the disappearance of condensate due to phosphorylation is reversible, we used CTD phosphatases to remove the phosphate groups and monitored condensation status. Ssu72/Symplekin phosphatase complex is a component of the 3’-end cleavage and polyadenylation factor (CPF) complex conserved throughout eukaryotes^26^. Its dephosphorylation activity is highly specific to phospho-Ser5 of the CTD heptad with no activity against phospho-Ser2^27,28^. While the Ssu72/Symplekin complex itself doesn’t form condensate (Figure S1F), we directly observed droplets’ appearance and gradual accumulation in pSer5 CTD sample upon its treatment with Ssu72/Symplekin (Figure 1D). The appearance of condensates coincides with the dephosphorylation process (Figure S1H). Control experiments using catalytically deficient phosphatase Ssu72 (C13D/D144N) exhibit no appearance of droplets under confocal microscopy, nor changes in absorbance at 350 nm (Figures 1D and 1E). Phosphatase-regulated CTD condensation was also observed for a second enzyme, human SCP1, which displays high dephosphorylation activity against both Ser2 and Ser5 of CTD heptad^29^. While purified human SCP1 phosphatase domain (residue 78-263) are homogenous in solution (Figure S1F), SCP1 treatment generates the condensate rapidly for both pSer5 CTD and pSer2 CTD (Figure 1F), corresponding with a sharp increase in 350nm absorbance in turbidity assay (Figure 1G) coinciding with dephosphorylation as confirmed by EMSA and MALDI-TOF analyses, (Figures S1I and S1J). Results from both phosphatase treatment experiments indicate that CTD phase transition caused by phosphorylation can be restored upon the removal of phosphate groups.

### Phospho-specific association of proteins with CTD promotes the reformation of droplets

In addition to phosphorylation, another well-established post-translational modification on the RNA polymerase II is prolyl isomerization^30,31^. Proline residues are frequently found in intrinsically disordered regions. Yet, it was unclear whether the cis/trans proline configuration influences LLPS behavior. Human prolyl isomerase 1 (PIN1) binds the CTD only when the serine of the Ser-Pro motifs in the heptad is phosphorylated – preferably at Ser5 but also at Ser2 albeit with weaker affinity and activity^32–34^. Therefore, we investigated how PIN1’s association with phospho-CTD affects condensate formation. Quantification of the PIN1 interaction with CTD peptide containing two heptad repeats with one Ser5 phosphorylation site estimate a K_d_ of 21 ± 7 µM while non-phosphorylated CTD does not bind, consistent with previous reports (Figure S2A)^31^. Neither PIN1 (Figure S2B) nor pSer5 CTD (Figure 1A) samples form droplets by themselves. When we added PIN1 into the pSer5 CTD solution, droplets appeared instantly (Figures 2A and 2B). The condensate increased with more PIN1, as visualized in confocal microscopy and turbidity assay, suggesting a dose-dependent effect (Figures 2A and 2B).

**Figure 2.**
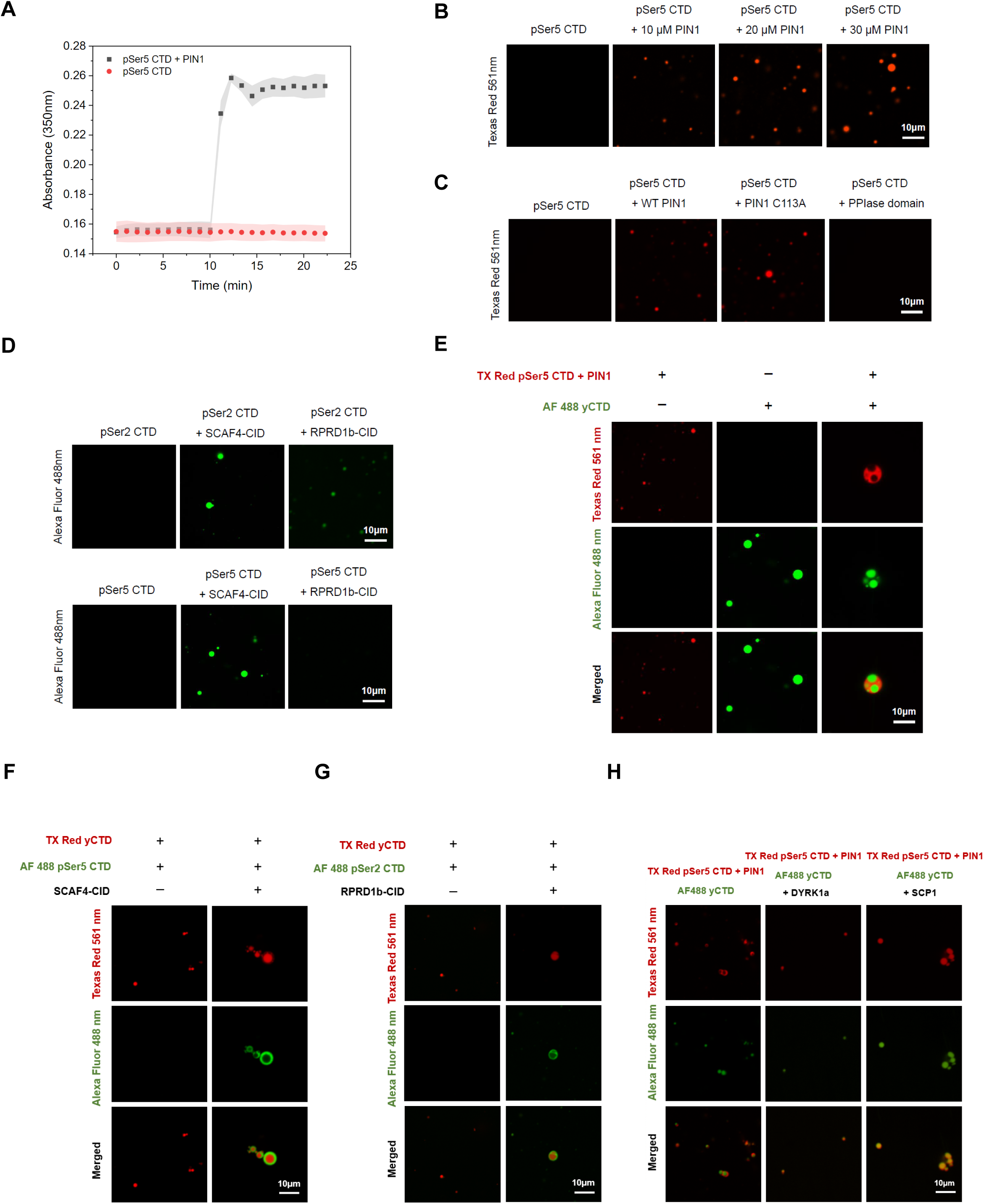
Phospho-specific association of proteins with CTD promotes the reformation of droplets, different CTD condensates remain distinct based on their physical properties. (A) Time-course turbidity assay of 10 μM pSer5 CTD samples treated with 10 μM PIN1 in the presence of 16% dextran. Addition of PIN1 at a specific timepoint (10 min) to the sample (black) was indicated with an arrow. The absorbance of control samples (pSer5 CTD only) is shown in red. Data points represent mean values of three replicate experiments, and error bars show the standard error. (B) Representative confocal microscopy images of 10 μM pSer5 CTD (Red, Texas Red-X) treated with different amounts of PIN1. (C) Representative confocal microscopy images of 10 μM pSer5 CTD (Red, Texas Red-X) treated with 10 μM of different PIN1 variants (Wild-type, C113A, PPIase domain). (D) Representative confocal microscopy images of 10 μM pSer2 CTD and pSer5 CTD (Green, Alexa Fluor 488) treated with 10 μM of different CTD binding proteins. (E) Confocal microscopy of unphosphorylated CTD (Green, Alexa Fluor 488) mixed with equimolar amount of pSer5 CTD (Red, Texas Red-X) and PIN1. (F) Confocal microscopy of unphosphorylated CTD (Red, Texas Red-X) mixed with equimolar amount of pSer5 CTD (Green, Alexa Fluor 488) and SCAF4. (G) Confocal microscopy of unphosphorylated CTD (Red, Texas Red-X) mixed with equimolar amount of pSer2 CTD (Green, Alexa Fluor 488) and RPRD1b. (H) Confocal image of immiscible mixture of unphosphorylated CTD (Green, Alexa Fluor 488) or pSer5 CTD bound to PIN1 (Red, Texas Red-X) incubated with kinase DYRK1a or phosphatase SCP1 overnight.

Two non-exclusive possibilities can explain the droplet-inducing effect of PIN1 on homogenously phosphorylated CTD. First, the enzymatic activity of proline isomerization by human PIN1 may alter the local conformation of the CTD backbone to induce LLPS. Alternatively, the interaction of PIN1 with phosphorylated CTD promotes condensate formation – potentially through a mechanism that neutralizes or shields the negatively charged phosphate groups. To distinguish the two possibilities, we utilized a catalytic-deficient PIN1 with the nucleophilic cysteine mutated to alanine, C113A^33^. When we added the PIN1 mutant to the phospho-CTD, we observed rapid droplet formation identical to the behavior seen for wild-type PIN1 (Figure 2C). On the other hand, when we preserved the PIN1 enzymatic domain but removed the substrate-recognizing WW domain (Figure S2D) ^31^, no condensate was observed (Figure 2C). Thus, the binding of PIN1 to phosphorylated CTD is necessary and sufficient to induce condensate formation. This observation raises questions about whether the recruitment of other phospho-CTD interacting factors could similarly counteract kinase-mediated CTD condensate dissolution, with implications for CTD condensate ‘switching’ or maturation as the CTD accumulates phosphates during the transcription cycle^11,13^.

The functional model of the CTD is to recruit different proteins to the RNA polymerase II at different stages of transcription based on its modification states^3^. Since the PIN1 association promotes condensate formation, we were curious if other proteins recruited to the phospho-CTD also induce phase separation through a similar binding-induced mechanism and investigated this directly. The CTD interacting domain (CID) is the largest CTD binding module recognized and found in many RNA binding proteins involved in splicing or termination^35^. To dissect only the CTD binding function, we isolated the CID domains from SCAF4. SCAF4 is reported as a CTD-binding protein binding to both Ser2 and Ser5 with a role in transcription termination^36^, whose binding are confirmed with phosphorylated CTD species using fluorescence polarization assay (Figures S2E). Mixing an increased concentration of SCAF4-CID with either pSer5 CTD or pSer2 CTD caused droplets to form visible with confocal microscopy and in turbidity assay (Figures 2D and S2G).

An intriguing inquiry arises regarding whether the condensate formation require a specific interaction between the phospho-CTD and the CID motif. To address this, we examined the purified CID domain of RPRD1b, an RNA-binding protein implicated in transcript elongation with specific binding to phospho-Ser2 but no association with phospho-Ser5 (Figures S2F)^37^. The introduction of increasing amount of RPRD1b CID to the pSer2 CTD resulted in an increasing in droplet formation, whereas the same experiment using pSer5 CTD remains homogenous (Figures 2D and S2H). This heightened concentration of RPRD1b induced more droplet scattering, as observed in turbidity assay (Figures S2H). Notably, the CTD binding proteins (PIN1, SCAF4, RPRD1b), purified to homogeneity in all these experiments, exhibited no phase separation under DIC microscopy (Figure S2B and S2I). Our experiments with CTD binding modules of recruited proteins (PIN1, SCAF4, RPRD1b) indicate that the association of phospho-CTD with binding partners enables condensate formation by the phosphorylated CTD.

### Different CTD condensates remain distinct based on their physical properties

Our biochemical results show that both unphosphorylated CTD and phospho-CTD are capable of condensation. This effect echoes the cellular observation that the initiation and elongation condensates coexist without fusion on the same gene loci^11^. To test if CTD and phospho-CTD condensates remain distinct or undergo fusion in vitro, we used unphosphorylated CTD labeled with Texas Red (emission at 561 nm) to mimic the promoter-bound RNA polymerase II and Alexa Fluor 488-labeled phospho-CTD (emission wavelength 488 nm) to imitate an active transcribing RNA polymerase II CTD. These labeled proteins were utilized in various mixing experiments in the presence or absence of CTD binding partner PIN1. Separately, unphosphorylated CTD forms red droplets in the presence of crowding agent dextran, and phospho-CTD forms green condensates with PIN1 mixed it, consistent with our previous observation (Figure 2E). Surprisingly, when we mixed the two CTD solutions in equal concentration, the two condensates remained immiscible, occasionally forming single condensates with a layered topology^38–40^ (Figure 2E). Prolonged incubation did not lead to the blending of the two condensates.

Observing distinct phases formed by unphosphorylated and PIN1-bound phospho-CTD prompted us to question whether this behavior is unique to PIN1. To test this, we mixed the unphosphorylated CTD with the phospho-CTD at a 1:1 ratio under the conditions in which Texas Red labeled unphosphorylated CTD phase separates while Alexa Fluor labeled pSer5 CTD remained homogenous (Figure 2F). The addition of the CID domains of SCAF4 induced droplet formation of the Alexa Fluor labeled pSer5 CTD, but no fusion between the unphosphorylated CTD and pSer5 CTD droplets was observed (Figure 2F). In some instances, the pSer5 CTD formed layered condensates with the unphosphorylated CTD (Figure 2F). Furthermore, we used a pSer2 CTD labeled with Alexa Fluor 488 to be mixed with unphosphorylated CTD (labeled with Texas Red). Adding pSer2 specific binding protein RPRD1b induced the green droplet formation but unphosphorylated CTD and pSer2 CTD droplets do not undergo fusion (Figure 2G). Collectively, these experiments indicate unphosphorylated CTD droplets do not mix with protein bound phosphorylated CTD droplets which coexist in distinct phases.

The biological implications are our results may be that RNA Pol II could be sorted between promotor associated CTD condensates and elongation associated phospho-CTD condensates as a function of phosphorylation state and factor association. Such a model requires the unphosphorylated CTD condensates to be both actionable substrates of CTD kinases and accessible to CTD interacting factors. Thus, we tested if we could induce condensate fusion by changing the phosphorylation status and binding interactions of the unphosphorylated and phosphorylated CTDs. We mixed Alexa Fluor 488 labeled unphosphorylated CTD, Texas Red labeled pSer5 CTD and phospho-CTD binding protein PIN1; as expected, the condensates remained separated (Figure 2H). However, upon incubation with the CTD kinase, DYRK1a, the previously unphosphorylated Alexa Fluor 488 labeled CTD colocalizes with the Texas Red labeled pSer5 CTD (Figure 2H). We next show the removal of CTD phosphorylation can induce similar fusion, by adding the CTD phosphatase SCP1. SCP1 dephosphorylates Texas Red labeled pSer5 CTD and enables its fusion with the Alex Fluor 488 labeled CTD condensates (Figure 2H). These results demonstrate that the PTM status of the CTD can dynamically dictate its partitioning between distinct condensate phases.

### CTD binding proteins colocalized with puncta formed by phosphorylated RNA polymerase II in cells

To corroborate our in vitro observation that association with CTD binding proteins can enable phospho-CTD condensate formation, we investigated the localization of CTD binding proteins relative to the phosphorylated RNA polymerase II in cells. We transfected YFP-labeled PIN1 into HeLa cells, which exhibited distinctive puncta formation localized to the nucleus (Figure 3A). The staining of RNA polymerase II with Ser5 phosphorylation also appeared as puncta, colocalizing exactly with PIN1 puncta (Figure 3A). When we transfected the IDR domain of MED1 to identify the possible location of Mediator^41^ (Figure S3A), the PIN1 puncta positions remained distinct from MED1 IDR domain puncta (Figure 3B), and the MED1 IDR didn’t colocalize with phosphorylated CTD (Figure S3B). To establish that the puncta we observed were not aggregates and remained dynamic in living cells, we employed fluorescence recovery after photobleaching (FRAP) beginning with MED1-IDR puncta whose dynamics are well established in the literature ^11,15,42^. Indeed, rapid FRAP on the timescale of seconds is consistent with liquid-like behavior observed for MED1-IDR (Figure 3C). Similarly, the puncta formed by PIN1 also recover at least partially after bleaching (Figure 3D), suggesting PIN1 partitions into and associates with the phospho-CTD condensates dynamically and is not aggregated. Compounded on our and others’ previous findings, these results indicate that the PIN1 phase separates specifically with phosphorylated RNA polymerase II.

**Figure 3.**
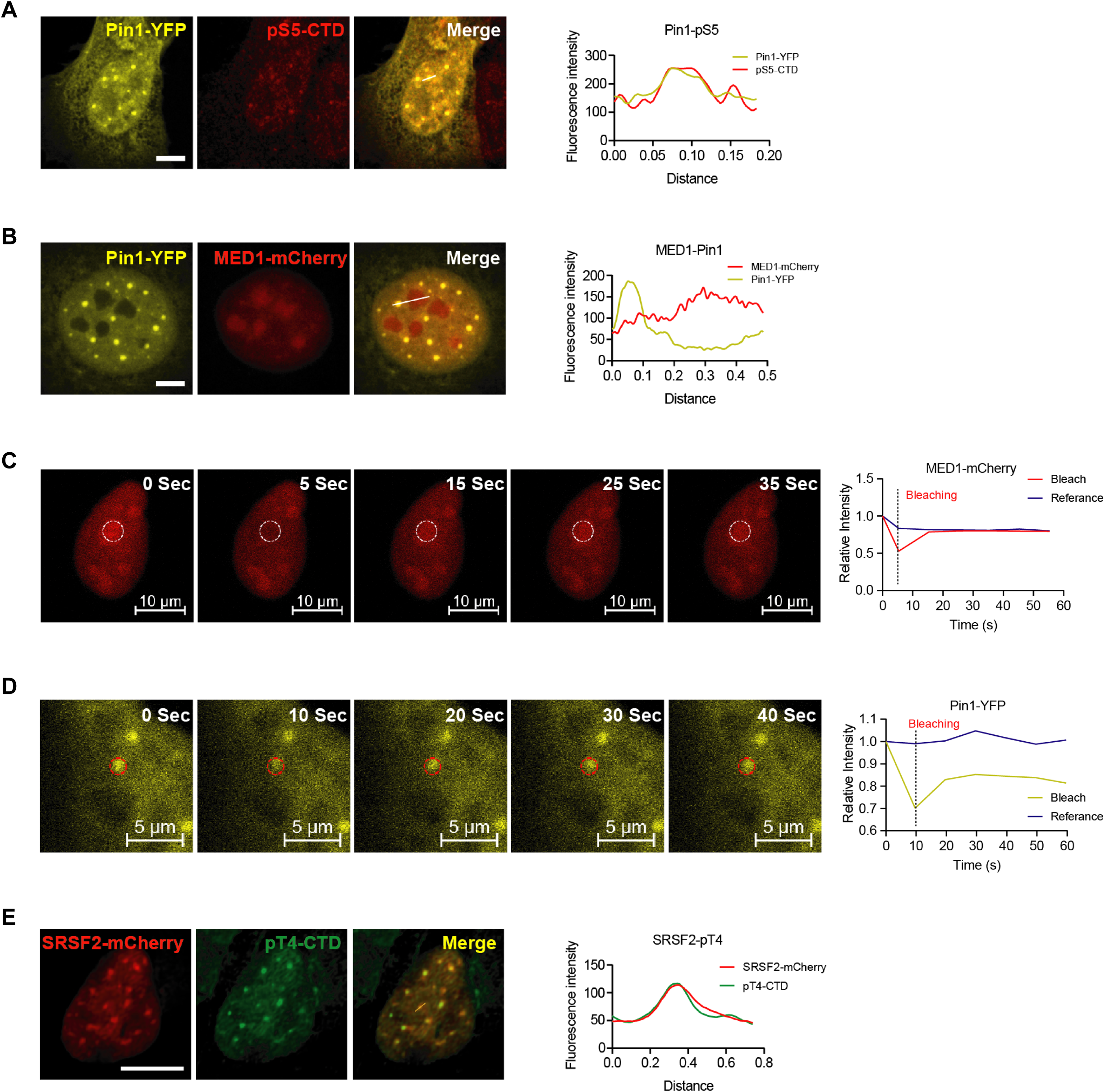
CTD binding proteins colocalized with puncta formed by phosphorylated RNA polymerase II in cells. (A) HeLa cells were transfected with PIN1-YFP construct and stained for phospho-Ser5 CTD antibody. Scale bars, 5 μm. (B) HeLa cells were transfected with PIN1-YFP and MED1 IDR-mCherry constructs, then fixed cells were detected by confocal microscopy. Scale bars, 5 μm. (C-D) U2OS cells were transfected with MED1 IDR-mCherry or PIN1-YFP constructs and fluorescence recovery after photo-bleaching (FRAP) assay to study the behavior of MED1 (C) or PIN1 (D) droplets in living cells. Scale bars, 10_μm (C) and 5_μm (D). Changes in the fluorescence intensity of droplets after photobleaching were plotted over time. (E) U2OS cells were transfected with SRSF2-mCherry construct and phospho-Thr4 CTD antibody. Scale bars, 5 μm.

To understand phosphorylated RNA polymerase II condensates in other biologically relevant contexts, we analyzed its localization with spliceosome components. Core spliceosome components, like SRSF2, were reported to partially form condensates with phosphorylated RNA polymerase II^11^. In cells, we found SRSF2 molecules in puncta with some overlapping significantly with phosphorylated Thr4 RNA polymerase II (Figure 3E), which also partial recover from photobleaching (Figure S3C). Overall, the images demonstrated that the puncta containing PIN1 and SRSF2 molecules colocalize with distinct Pol II harboring distinct patterns of phosphorylation and possess dynamics consistent with liquid-like, not aggregate, behavior. These observations support a model of compositionally distinct Pol II condensates, governed at least in part by the phospho-specific association of CTD interacting factors.

### Effect of electrostatics charges on CTD condensates

To link our biochemical observation to transcriptional function, we engineered the consensus CTD to mimic the phosphorylation state at different sites of the heptad. We hypothesize that reversible phosphorylation of the CTD dissolves the formed droplet because the repulsion between negatively charged CTD molecules counteracts attractive interactions, such as π-π interactions^43^. If this hypothesis holds, a negative charge installed at any position within each CTD heptad would disrupt condensate formation. To test this hypothesis, we employed phosphomimetic mutations, inserting glutamate at T4, S5, or S7 position of each heptad (Figure S1A) and compared the condensation of each variant in vitro (Figures 4B, S4A and S4B). Unlike the wild-type CTD (Figure 4A), no condensation was observed even in the concentrations of protein greatly exceeding the saturation concentration for the wild type CTD or with elevated concentrations of crowding agent (Figures 4B, S4A and S4B), consistent with turbidity assay showing no absorbance for light scattering by droplets (Figure S4D).

**Figure 4.**
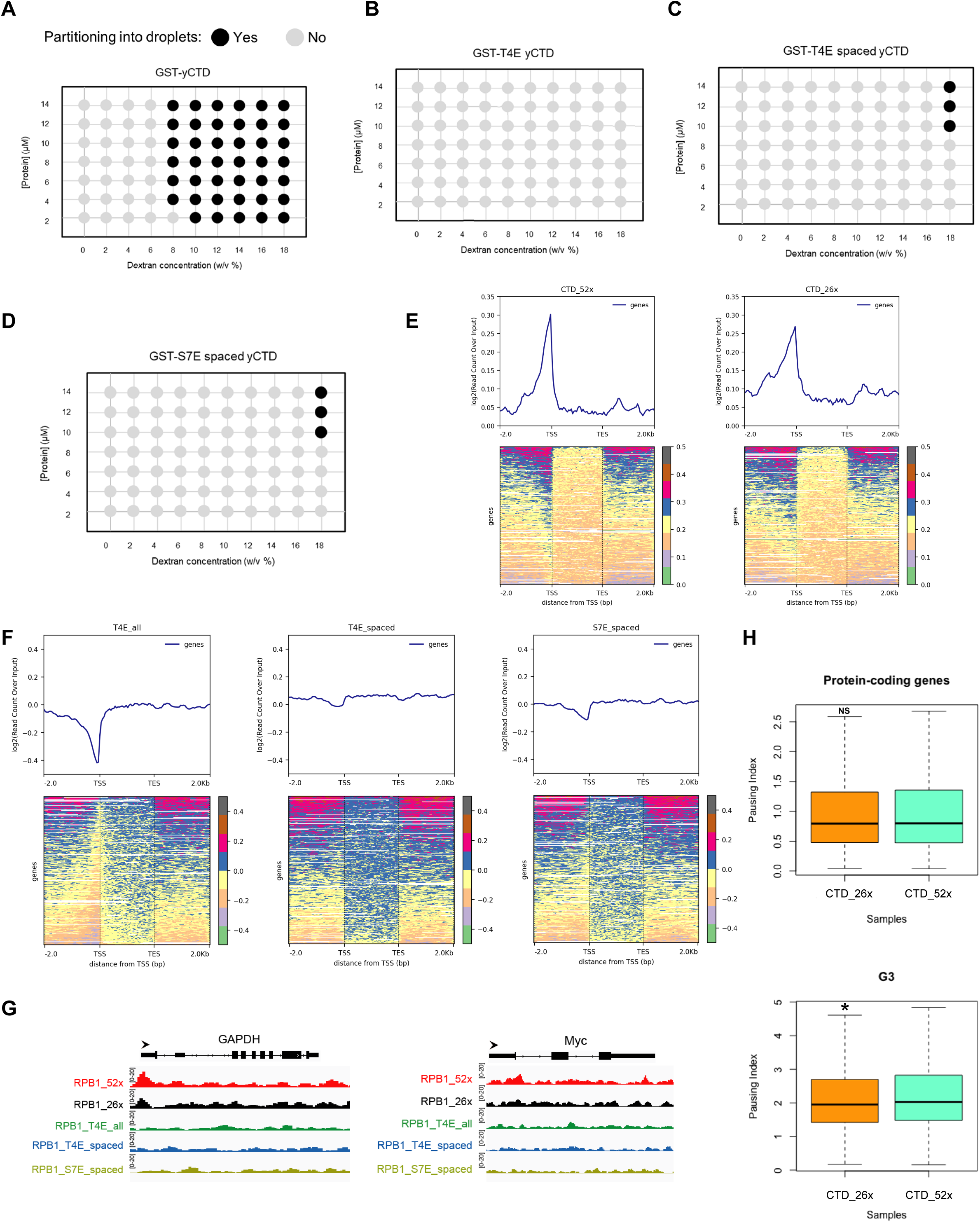
CTD condensation properties in vitro predict genomic locations of RNA polymerase II in vivo. (A) Phase diagram of wild-type GST-yCTD. Black filled circles indicate conditions where liquid-liquid phase separation is observed with DIC microscope. (B) Phase diagram of GST-T4E yCTD. (C) Phase diagram of GST-T4E spaced yCTD. (D) Phase diagram of GST-S7E spaced yCTD. (E) ChIP-Seq analyses on the distribution of RPB1_52xCTD and RPB1_26xCTD in the genome. Average peak coverage is shown in a bin size of 50 bp for a window 2 kb upstream from TSS and 2 kb downstream of TES. (F) ChIP-Seq analyses on the distribution of RPB1_26xCTD T4E (all), 26xCTD T4E (spaced) and 26xCTD S7E (spaced) in the genome. (G) ChIP-Seq example illustrating the association of different RPB1 CTD constructs along with the active transcribing genes. (H) Boxplots on the pausing index changes from all cluster of genes (19630 genes) and G3 (4908 genes) clusters.

We then asked whether the frequency and spacing of the phosphomimetic negative charge affected phase separation. To test that, we mutated every other T4 or S7 to negatively charged glutamate residues (T4E-spaced or S7E-spaced) (Figure S1A). These spaced variants can form liquid droplets (Figures 4C and 4D), but the concentration of protein and dextran needed for the liquid phase to appear is much higher than that of the wild-type CTD. The effect of negative charges was dose dependent ― the wild-type CTD (no negative charge) (Figure 4A), T4E-spaced and S7E-spaced (one negative charge every two heptads) (Figures 4C and 4D), S5E, T4E, and S7E (one negative charge every heptad) (Figures 4B, S4A and S4B) display a correlation between the density of negative charges to the loss of phase separation, regardless of their position. These results suggest that threshold levels of negative charge impair CTD condensation and provide an experimental platform to relate CTD condensation properties in vitro with functional effects on RNA polymerase II transcription in vivo.

### CTD condensation properties in vitro predict genomic locations of RNA polymerase II in vivo

The CTD of RNA polymerase II is crucial to eukaryotic transcription, yet this variation’s functional and evolutionary relevance remains unclear. To study the effects of CTD condensation in the context of the full polymerase, we introduced RPB1 plasmids harboring 52xCTD (human wild-type), 26xCTD consensus heptads, and 26xCTD mutants heptads with YFP in HEK 293T cells. Previously, multiple labs have shown that 52xCTD is more disposed to condensate formation than 26xCTD^10,44^. For 26xCTD mutants, we focused on T4E/S7E mutants to dissect the effects of CTD charges on transcription without the confounding variable of S2 and S5 phosphomimetic mutations that abolish transcription^45^. We performed Chromatin Immunoprecipitation (ChIP)-Seq analysis to identify genomic locations of Pol II harboring the CTD mutants with differing condensation properties. We first validated the function of the ectopically expressed Pol II mutants in HEK 293T cells (Figures S4E and S4F), then preformed ChIP-Seq. Each Pol II mutant was immunoprecipitated using anti-GFP antibodies to specifically map distribution of the mutant Pol II isoforms. Although the transcription start site (TSS) associated peak in 26xCTD profile was slightly lower than in 52xCTD, the profiles of RPB1_52xCTD and 26xCTD were very similar (Figure 4E), consistent with a role for CTD condensation in targeting Pol II to promoters^10,44^. Intriguingly, the binding of RPB1_26xCTD T4E/S7E mutant was completely lost genome-wide, most notably in the TSS region (Figure 4F). The distribution of RNA polymerase II along the gene body was altered exemplified in multiple genes (Figure 4G). The degree of global impairment of Pol II distribution as a function of negative charge is consistent with a model of impaired phospho-Pol II recruitment to unphosphorylated Pol II condensates predicted by our in vitro results.

To quantify the change of distribution of the polymerase over gene regions, we calculated the “Pausing Index” (PI) as the ratio of Pol II read density near the promoter (–50 to +300 bp of Transcription Starting Site Region/TSSR) over the remainder of the gene body (+300 downstream of the TSS to +3 kb past the Transcription Termination Site/TTS) ^46^. The protein-coding genes (n =19,630) were clustered into four groups based on the pausing index (Figure S4G). Comparison of the pausing index of all the genes showed no statistical difference between 26xCTD and 52xCTD, but slightly decreased PI (*p* = 1.611e-06) was observed in G3 cluster of most-paused genes in 26xCTD (Figure 4H), suggesting 26xCTD does not substantially alter Pol II pausing behavior globally, consistent with previous report^47^. Taken together with our biochemical data (Figures 4A), these results showed both 52xCTD and 26xCTD are competent to partition into unphosphorylated Pol II clusters at the promoters genome wide. Conversely negatively-charged phospho-CTD Pol II mutants are impaired in their ability to partition into unphosphorylated Pol II condensates resulting in failed initiation that scales with negative charge.

### Growth study and transcriptomic analysis of RNA polymerase II with different CTD variants

To interrogate the role of different CTD negative charge mutants in vivo without the confounding variable of endogenous wild-type Pol II, we generated YFP_RPB1_52xCTD, 26xCTD and 26xCTD T4E/S7E constructs in the context of a mutant version of RPB1, N792D, conferring resistance to the potent Pol II inhibitor, α-amanitin^48^. Expression of the α-amanitin resistance Pol II mutants was similar, measured by both fluorescence intensity and western blotting (Figures 5A-5B and S5A-S5B). After 72 h of α-amanitin administration to degrade endogenous Pol II, we evaluated the physiological function of transfected Pol II mutants. The results showed that both RPB1_52xCTD and 26xCTD could sustain cell viability, with the RPB1_26xCTD exhibiting slowed growth^47^ (Figure 5C). The T4E/S7E mutants exhibited a greater reduction of cell survival after endogenous RPB1 depletion by α-amanitin (Figures 5A and S5A), consistent with impaired CTD condensation in vitro and Pol II promoter association in cells (Figure 4B-4D and 4F).

**Figure 5.**
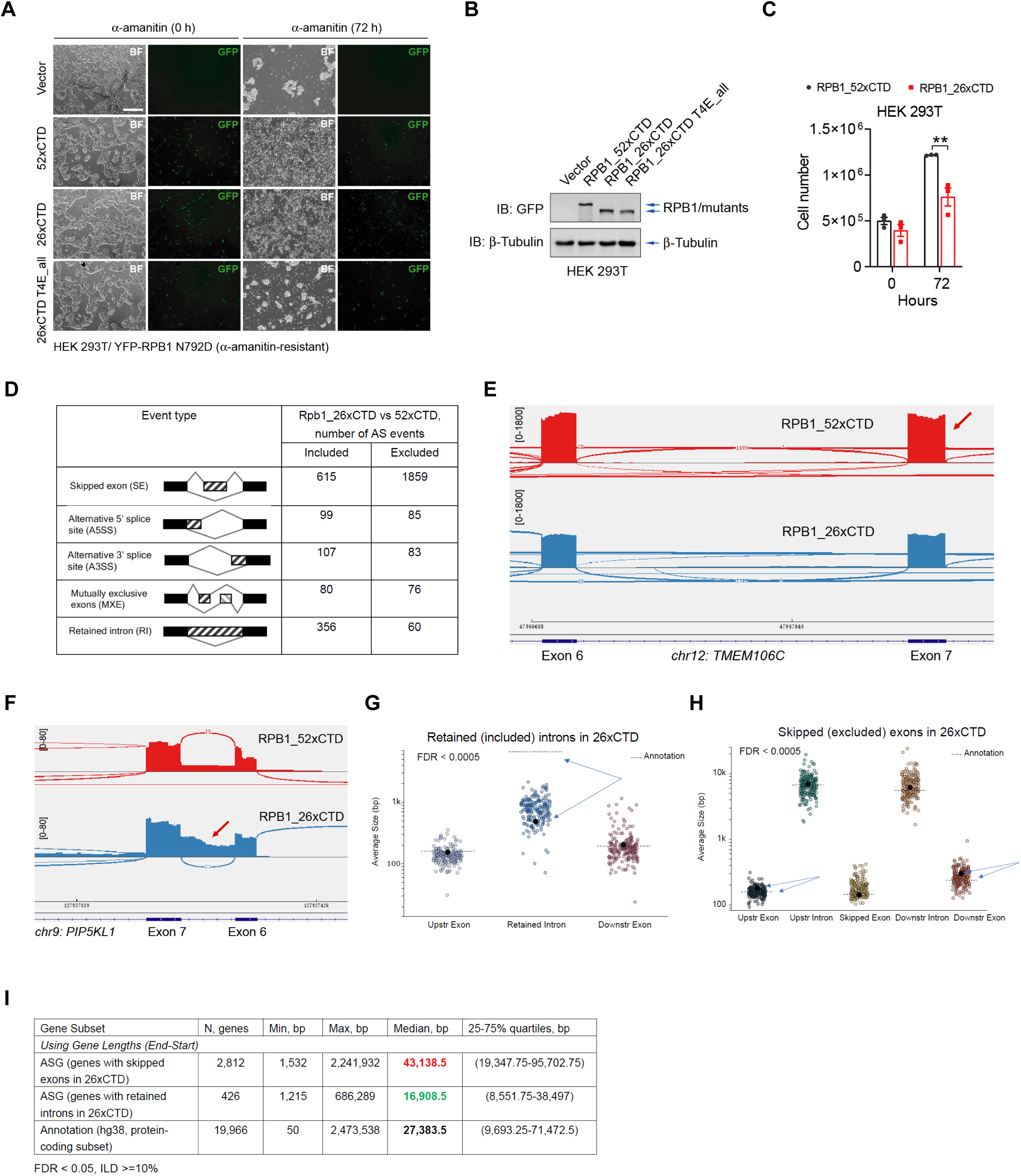
Different CTD variants alter cell survival and alternative splicing. (A) Cell growth of HEK 293T which transfected with RPB1_52xCTD, 26xCTD and 26xCTD T4E (all) constructs upon α-amanitin administration. (B) Western blotting detected the expression level of RPB1_52xCTD, 26xCTD and 26xCTD T4E (all) constructs. (C) Cell density in RPB1_52xCTD and 26xCTD group upon α-amanitin administration. (D) Types and absolute numbers of annotated alternative splicing events (ASE) that were significantly different in RPB1_26xCTD vs. RPB1_52xCTD cells (with parameters FDR < 0.0005). In event type illustration, the constitutive exon is black, whereas alternatively, spliced exons are striped. (E-F) Examples of Sashimi plots of *TMEM106C* and *PIP5KL1* genes in RPB1_26xCTD compared to RPB1_52xCTD: read densities are shown on the y-axis. Arrow indicates the inclusion of an alternative exon or intron. (G-H) Average intron (G) or exon (H) sizes for potential alternative splicing events in RPB1_26xCTD, derived from input annotation file which indicated by the dashed ‘Annotated’ line. Only high-confidence AS events (FDR < 0.0005) are calculated in plots. (I) Calculated the lengths of genes with skipped exons and retained introns in RPB1_26xCTD compared with annotation (with parameters FDR < 0.05; ILD, inclusion level difference, ≥ 10%).

To further dissect the role of different CTD repeats in transcription and cell viability, we performed whole-transcriptome RNA-Seq to detect the polyadenylated mRNA in RPB1 N792D mutant with 52 or 26xCTD repeats, respectively. Differential expression analysis (Figure S5C) in RPB1_26xCTD compared to 52xCTD revealed that among a total of 39,146 annotated expressed genes (counts > 0), there were only 675 genes upregulated (1.7%) and 742 genes (1.9%) downregulated in 26xCTD (log_2_FC cutoff = 0.58, *p*-adjusted cutoff = 0.05). This data suggests that 26xCTD only modestly alters transcription globally. Additionally, analyzing occupancy data in ‘DiffBind’ R package, we derived and overlapped 26xCTD and 52xCTD consensus ChIP-Seq peaks, as well as their shared peakset, with genomic locations of putative enhancers in HEK 293T cells from EnhancerAtlas annotation^49^. As a result, nearly half of the shared peaks overlapped enhancer regions, and a total percentage of peaks overlapping enhancers was higher in 52xCTD (21.5%) compared to 26xCTD (15.4%) (Figure S5D). These results are consistent with the previous observation that CTD shortening impairs enhancer transcription^47^.

### The length of CTD alters alternative splicing

Previously, the observation of Pol II initiation and elongation condensates have been reported^11,13^. Our biochemical and cellular studies suggested that Pol II with impaired condensation could be associated with splicing defects. The truncated 26 heptad CTD brings to mind the length of CTD in *S. cerevisiae*, where splicing events are rare compared to human (52 heptad CTD). Using RPB1_26xCTD, which supports viability despite condensation defects, we investigated the splicing outcome of the truncated CTD. To test this, we used rMATS-turbo^50^ followed by SpliceTools^51^ suite to analyze splicing differences in shortened (26x) vs. full-length (52x) CTD. We identified substantial differences with a total number of 16,168 significant alternative splicing (AS) events in RPB1_26xCTD vs. RPB1_52xCTD (FDR < 0.05) (Figure S5E). We set an additional constraint [ILD>=10%], where ILD (inclusion level difference) is an index of the strength of splicing events. As a result, a total of 9,239 AS events were found statistically different between conditions (FDR < 0.05, ILD >=10%), with significantly more exon skipping (SE) (4,350 events in 2,812 unique genes) and intron retention (RI) (511 events in 426 unique genes), being 1.9-fold and 2.9-fold more frequent in 26xCTD, respectively (Figure S5F). The difference is more significant when we focused on high-confidence AS events (FDR < 0.0005), in which SE frequency was 3 fold and RI frequency 6-fold for RPB1_26xCTD vs. RPB1_52xCTD (Figure 5D). All data indicate that 26xCTD leads to more frequent exon skipping and intron retention. Specific examples of SE (ILD = 100%, FDR =2.95e-06) (Figure 5E) and RI (ILD =51.4%, FDR =1.8911e-10) (Figure 5F) events in RPB1_26xCTD condition vs. RPB1_52xCTD are shown as Sashimi plots. Using the SpliceTools suite, we found that the global footprint of AS on the cell transcriptome of the 26xCTD cell resembles that of the knockdown of spliceosome components (Figure S5G). There was a significant level of increased exon skipping in 9% (fraction = 0.0879) of all expressed genes with TPM >= 3, which was similar to the effect of SRFBP1 (RNA-binding protein/RBP) knockdown but somewhat lower than the effects of the core spliceosome knockdown (U2AF1 and U2AF2)^51^ (Figure S5G).

Further analysis of the skipped exon/retained intron sizes in 26xCTD condition showed that the average size of retained introns upon 26xCTD was considerably shorter than the ones from the global average derived from an input annotation file (Figure 5G). On the contrary, exons located upstream/downstream from the skipped exons in 26xCTD were slightly longer than the annotation mean (Figure 5H). Consistent with this, analysis of the length of the genes with SE or RI in 26xCTD revealed that exon skipping events were associated with longer genes and intron retention was associated with shorter gene lengths compared to hg38 annotation median (Figure 5I). The analysis with 19,966 unique protein-coding genes suggest that 26xCTD has reduced accuracy when including exons in longer genes and/or cutting out introns from shorter genes. The binding of U1 splice donor factors and U2 splice acceptor factors to initial transcripts depend on the quality of the splice junction sequences. Using SE/RI SpliceSiteScoring, we assessed the quality of all splice junction sequences for significantly altered SE and RI events in RPB1_26xCTD, which showed that short CTD facilitated skipping of exons with weak splice site scores (Figure S5H) and retaining introns that are surrounded by “weaker-scored” donor and acceptor exons (Figure S5I). Taken together, these data imply that the short CTD likely have defects in spliceosome recruitment to RNA Pol II condensates leading to aberrant splicing.

To better understand the physiological impact of impaired splicing in the context of CTD mutations, we used ‘SETranslateNMD’ from the SpliceTools suite to understand the consequences of exon skipping caused by 26xCTD. Intriguingly, ∼43% of alternative transcripts associated with a high-confidence set of skipped exons by Pol II 26xCTD were predicted to undergo nonsense-mediated mRNA decay (NMD) (Figure 6A). Therefore, 26xCTD-mediated exon skipping can frequently lead to NMD of transcripts, decreasing their effective expression level (Figure 6B). Overrepresentation analysis showed that frameshifted transcripts were enriched for genes playing a role in cell cycle transition (Figure 6B), which was consistent with decreased proliferation of 26xCTD transfected cells. Some representative genes with exon skipping under 26xCTD, *AURKB* (ILD = −0.142, FDR = 1.56e-11) and *CDK4* (ILD = −0.026, FDR = 3.96e-0.5) are demonstrated as examples of NMD-transcripts (Figures 6C and 6D). These genes are known as master regulators of the cell cycle and DNA replication^52,53^, and previous report also showed alternative splicing governs cell cycle progression through NMD genes, including AURKB^54^.

**Figure 6.**
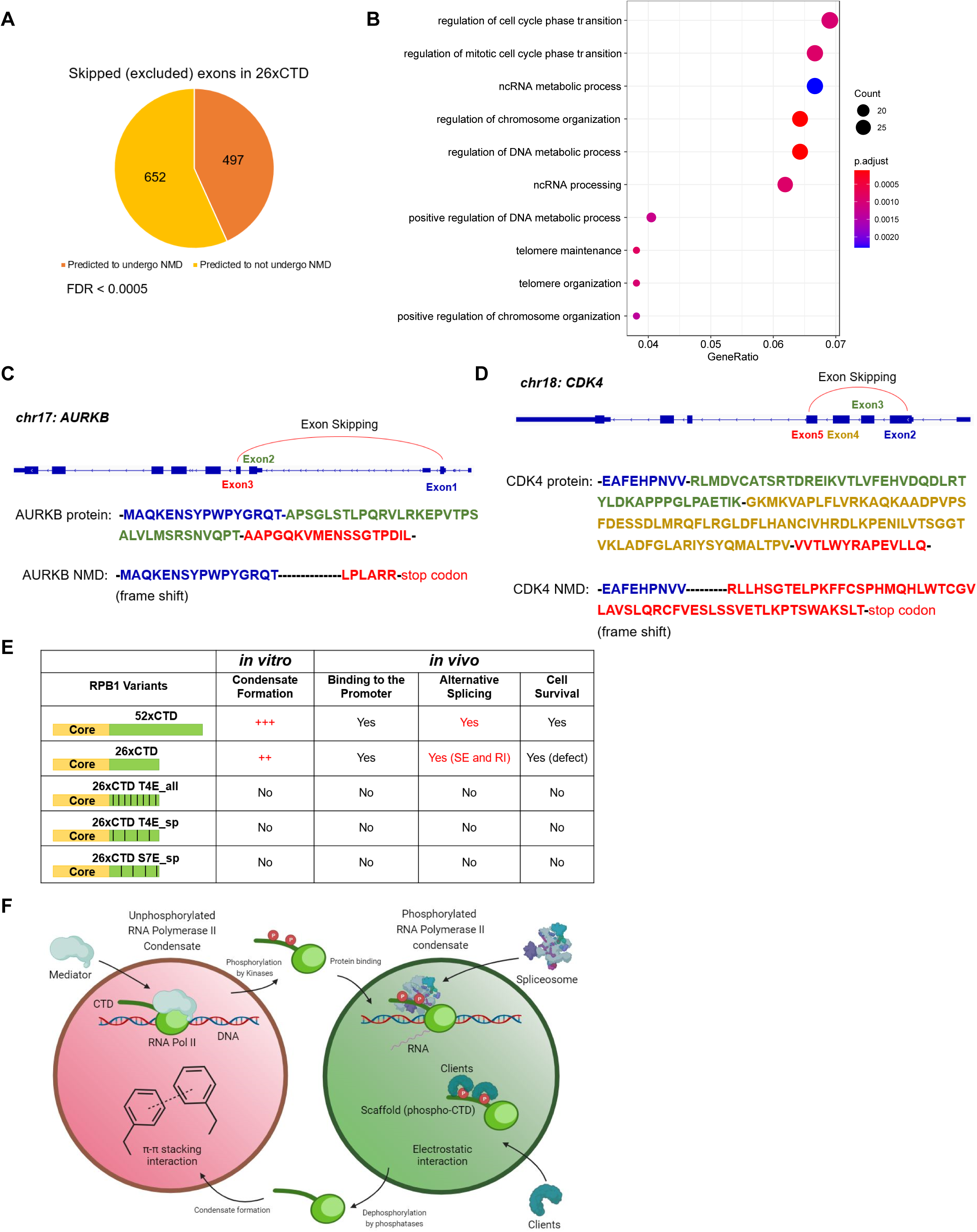
The length of CTD governs cell growth by the regulation of NMD (nonsense-mediated mRNA decay) of transcripts. (A) NMD or non-NMD of transcripts detected in skipped exons in RPB1_26xCTD (with parameters FDR < 0.0005). (B) Gene ontology of biological processes enriched among NMD (nonsense-mediated mRNA decay) of genes in RPB1_26xCTD skipping exons. (C-D) Examples of genes related to cell cycle in RPB1_26xCTD skipping exons, leading to frame shift and premature termination of gene transcription. (E) Comparison of the ability in condensate formation in vitro and biological function in vivo among multiple CTD variants. (F) Model for phase separation sorting of RNA polymerase II.

Overall, these results support our hypothesis that the length of CTD affects functional transcriptional condensate assembly, is involved in the recruitment of splicing factors, and can lead to defects in recognizing precise splice sites in case of shortened CTD repeats.

## Discussion

It was well accepted that a primary function of CTD was to facilitate pre-mRNA splicing by recruiting splicing factors^20^. However, recent high-resolution cryo-EM structures of spliceosomes have revealed no observable physical interaction with the CTD^30,55–58^. In this study, we provide a molecular mechanism for this apparent discrepancy. The interaction between phospho-CTD and splicing factors promotes the formation of splicing condensates, increasing the local concentration of splicing components. This, in turn, facilitates the spliceosome assembly and enhances the efficiency of pre-mRNA processing.

Through cellular imaging, we have observed that phosphorylated RNA polymerase II associates with spliceosome components, forming distinct puncta. These puncta exhibit liquid-like dynamics. Interestingly, the expansion of the CTD length during evolution seems to align with the increased frequency and complexity of splicing events in eukaryotic cells. Fine-tuning the capacity for condensate formation inside cells appears to play a crucial role in splicing precision. Notably, when we examined an RPB1 variant with shortened CTD comprising 26 heptad repeats, we observed significant differences in alternative splicing patterns (Figure 6E). This result differs from a previous study that little difference was found when only long transcripts were sequenced^47^. In our study, the most significant difference lies in the short genes when the introns end up included up to six-fold more frequently. In longer genes, we saw a tendency of exon inappropriately skipped. Such errors were more pronounced at weaker splicing sites. We highlight that truncation of the CTD by a factor of two impairs splicing outcomes by a factor of 3 for skipped exons and 6 for retained introns. One implication of this non-linear impairment is that altered CTD phase separation, not simply a reduction of splicing factor binding sites by half contributes to splicing defects for Pol II mutants with truncated CTDs consistent with the ability of phase transitions to transform linear inputs (CTD length) into non-linear functional outputs with consequences for organismal fitness (splicing)^59^. Intriguingly, these alternative splicing events affect numerous genes that regulate cell growth. The alternatively spliced isoforms encoded shorter proteins with premature termination, providing an additional explanation for the observed deficiency in cellular growth associated with shortened CTD beyond impaired promotor recruitment.

In our study, we generated multiple RPB1 variants with negative charges and consistently observed their inability to bind to the promoter for transcription initiation (Figure 6E). This finding aligns with previous research indicating that promoter binding requires unphosphorylated RNA polymerase II^60^. The assembly of the PIC requires hydrophobic interaction facilitated by unphosphorylated CTD (Figure 6F). The recently published structure of Mediator and RNA Pol II^61^ revealed that a residual CTD fragment bridges the interface between the Middle and Head of Mediator complex involving Y1 residues of CTD heptad making hydrophobic contacts^62–64^. The high local concentration of the Pol II at the site of transcription initiation can account for the phenomenon of transcriptional bursts. While dominated by hydrophobic forces like π-π stacking as seen in unphosphorylated CTD and Mediator, once transcription starts, CTD kinase-mediated phosphorylation of Pol II disrupts these hydrophobic interactions (Figure 6F). As phosphorylation accumulates on the CTD, it recruits phospho-specific binding proteins, predominantly RNA-processing factors (Figure 6F). In this scenario, phospho-CTD functions as the scaffold for multivalent interaction with the transcription regulatory proteins (clients), evolving the initiation condensate into one tuned for splicing. The existence condensates with layered topologies in other membraneless organelles^40^. For example, nucleolus contains subcompartments representing distinct coexisting condensates^65^. The layers of condensates give rise to a rational, organized RNA synthesis and folding/processing factory as the RNA matures. Similarly, layers of condensates have been observed in coexisting chromatin condensates^66^. Our in vitro and in vivo study of RNA polymerase II suggests a model where transcription initiation condensates including unphosphorylated CTD evolve to transcript-processing condensates as a function of Pol II phosphorylation and splicing factor recruitment. Our results shed new light of LLPS into the classic model of CTD function^20^ where phosphorylation acts to recruit transcriptional regulators not only via binding but by recruitment to functional condensates, highlighting how the CTD variations with different physical properties can affect splicing outcome perhaps by altering the local concentration of splicing factor clients of Pol II condensates.

## Acknowledgments

We thank Dr. Jack E Dixon and Tanja Mittag for advice on the manuscript and the National Institutes of Health (R01GM104896, R01GM125882 and R35GM148356 to YJZ) for supporting our research. Thanks also to technical assistance by the core facilities of the University of Texas at Austin. The content is solely the responsibility of the authors and does not necessarily represent the official views of the National Institutes of Health.

## Author contributions

Q.Z. and W.K. carried out most experiments and helped with experimental design. S.P. conducted the bioinformatics analysis, J.M. and B.P. helped with experimental design and discussions. Y.J. Z. conceived the study and experimental design, Y.J. Z., Q.Z., W.K. and B.P. wrote the manuscript.

## Methods

### Bacterial strains, Cell lines, Reagents, and Antibodies

All *E.coli* strains were grown in L.B. (Luria-Bertani) or Terrific Broth media at 37°C as indicated below. HeLa, HEK 293T and U2OS cells were from ATCC. No cell lines used in this study were found in the database of commonly misidentified cell lines maintained by ICLAC and NCBI Biosample. All cell lines were cultured in DMEM medium with 10% fetal bovine serum (FBS) and 1% penicillin-streptomycin solution at 37°C in 5% CO_2_ (v/v).

Polyethyleneimine (PEI, Polysciences) and FuGENE HD (Promega) transfection reagents were used according to manual instructions and transfected plasmids in cultured cells. The α-amanitin (sigma) was used for endogenous RPB1 degradation, the concentration is 2.5 μg/mL.

The monoclonal antibodies anti-pSer5 CTD S2 (Millipore, 04-1571, 1:500 dilution), anti-pSer5 CTD S5 (Millipore, 04-1572, 1:500 dilution), the monoclonal antibody anti-pSer5 CTD T4 (Active Motif, 61361, 1:200 dilution). The anti-GFP (Proteintech, 50430-2-AP) was used for Chip experiment 5 μg/sample. The anti-rabbit IgG antibody was purchased from Invitrogen (08-6199).

### Constructs

All 26x yCTD constructs (WT, S5E, S7E, S7E in every other repeat, T4E, T4E in every other repeat) were ordered as synthetic genes (Genscript), amplified, and cloned using ligation-independent cloning (SLIC)^33^. For bacterial protein expression, all constructs were cloned into a PET28a vector (Novagene) containing a 6x histidine tag and glutathione-S-transferase (GST) tag with a 3C protease cleavage site added after the two tags. For mammalian protein expression, YFP-RPB1-WT (52xCTD) was obtained from Addgene, YFP-RPB1-26xCTD and T4E/S7E mutants were generated by PCR-based cloning performed by a kit from ThermoFisher. For HeLa cell transfection & imaging, PIN1 were cloned into a pcDNA3 vector containing YFP tag with a 3C protease cleavage site. MED1 IDR was kindly shared from Dr. Jason Liu’s lab in University of Texas Health Science Center at San Antonio and containing mCherry tag. SRSF2 was obtained from Addgene. All coding sequences were verified by DNA sequencing.

### Protein Purification

*E.coli* (DE3) cells were used as the protein expression system for the proteins used in the study. The transformation was carried out by thawing the competent cells on ice for 5 min, adding the DNA to cells and incubating on ice for 30 min, heat shocking at 42°C for 90 seconds, and finally cooling the cells on ice for 3 min. The cells were recovered in SOC medium for 1 h at 37°C and were plated on Luria-Bertani agar plates containing 50 μg/mL kanamycin for selection. Individual colonies were grown in 50 mL of Luria-Bertani medium at 37°C containing 50 μg/mL kanamycin. 1 L of terrific broth medium (Thermo Fisher) was inoculated with 10 mL of inoculum, and the culture was grown to an O.D. of 0.4-0.6. 0.5 mM IPTG was added to each culture to induce the protein expression. The cultures were pelleted by centrifugation after overnight growth (20 h at 16°C), and the cells were lysed through sonication in a lysis buffer (50 mM Tris-Cl pH 8.0, 500 mM NaCl, 10% glycerol, 0.1% Triton-X 100, 20 mM imidazole, and 10 mM BME). Sonication of cell pellets were carried out on ice at 90 A for 3 min per cycle (1 s on and 5 s off) for five cycles with a 3 min break between each cycle. The lysate was cleared by centrifugation at 27000 g for 45 min at 4°C. The supernatant was purified through affinity column chromatography using Ni^2+^/NTA beads (Qiagen). The column was equilibrated with lysis buffer. Then, the cleared lysate supernatant was run through the column. The column was washed with 10 times column volume of wash buffer (50 mM Tris-Cl pH 8.0, 500 mM NaCl, 10% glycerol, 20 mM imidazole, and 10 mM BME) and eluted with an elution buffer (50 mM Tris-Cl pH 8.0, 500 mM NaCl, 10% glycerol, 250 mM imidazole, and 10 mM BME). Proteins were dialyzed in a gel filtration buffer (50 mM Tris-Cl pH 8.0, 500 mM NaCl, 10 mM BME) at 4°C overnight. Proteins were concentrated using centrifugal concentrator (Sartorius) and further purified with size exclusion chromatography using a Superdex 200 column (GE Life Sciences). The purity of each protein fractions was assessed by polyacrylamide gel electrophoresis (Coomassie Brilliant Blue Staining).

### Covalent labeling of the CTD molecules

Two different kinds of succinimidyl ester probe (Invitrogen) that contain different fluorescent dye (Texas Red^TM^ and Alexa Flour 488) were purchased and stored at −20°C as powder samples. 1 mg of each dye was dissolved in 100 μL of DMSO and mixed with 1 mL of 10 mg/mL GST-yCTD protein sample. Each protein-dye mixture was incubated for 1 hour at 25°C with continuous stirring. Then the protein-dye conjugate was separated from the unreacted dye by using size-exclusion chromatography. A Superdex 200 column (GE Life sciences) was equilibrated with PBS, and the reaction mixture was separated by using PBS as the gel filtration buffer. The fraction corresponding to conjugated GST-yCTD were collected and concentrated for storage at −80°C.

### CTD phosphorylation

GST-yCTD was phosphorylated using either *h.sapiens* ERK2 or GST-tagged *h.sapiens* DYRK1a. 5 mg/mL GST-yCTD were incubated with 0.125 mg/mL ERK2 or 0.5 mg/mL GST-DYRK1a, supplemented with 50 mM TRIS pH 7.5, 5 mM ATP, and 5 mM MgCl_2_. After overnight incubation at 30°C, the phosphorylation reaction was quenched by adding EDTA to a final concentration of 5 mM. After completion of overnight reaction, phosphorylated GST-yCTD was mixed at a final concentration of 10 μM into 16% dextran, 50 mM TRIS (pH 7.5), 150 mM NaCl, 10% glycerol, and 1 mM DTT in order to check condensate formation with microscopy or turbidity assay (as described below).

### Kinase and phosphatase treatment assay on condensates

Before kinase treatment, GST-yCTD droplets were formed by mixing 10μM GST-yCTD in 50 mM TRIS (pH 7.5), 150 mM NaCl, 10% glycerol, 1 mM DTT with 16% dextran. After droplet formation, 2 mM ATP, 2 mM MgCl_2_, and appropriate amount of kinase (ERK2 or DYRK1a) were mixed and incubated at 30°C with continuous shaking. For phosphatase treatment experiments, phosphorylated GST-yCTD sample was mixed at a final concentration of 10 μM into 16% dextran, 50 mM TRIS (pH 7.5), 150mM NaCl, 10% glycerol, and 1 mM DTT, then appropriate amount of phosphatase (SCP1 or Ssu72) was added and incubated at 30°C with continuous shaking. Droplet disruption or formation was monitored by microscopy or turbidity assay (as described below). Phosphorylation of GST-yCTD samples was confirmed by using gel shift assay (EMSA) and MALDI-TOF mass spectrometry (as described below).

### Turbidity assay

Turbidity assays were carried out in 50 μL samples containing 50 mM TRIS (pH 7.5), 150 mM NaCl, 10% glycerol, 1 mM DTT, indicated concentrations of GST-yCTD or GST-yCTD variants, and indicated concentration of dextran. Each solution was prepared in 96 well plate (Thermo Scientific) and absorbance at 350 nm readings were taken in a plate reader (Tecan) using default Absorbance settings. For kinase and phosphatase treatment assay, absorbance at 350 nm was measured at 30°C with continuous shaking.

### MALDI-TOF Mass Spectrometry and EMSA

5 μL of phosphorylated GST-yCTD samples were taken out from reaction batch for measuring molecular weight. The samples were desalted over Ziptip C18 resins (MilliporeSigma) using standard protocols. Mass spectrometric analysis of phosphorylated GST-yCTD was carried out in an AB Voyager-DE PRO MALDI-TOF (Brunker Corporation) with the 1:1 DHB matrix (Thermo Fisher Scientific).

GST-yCTD samples treated with kinases were analyzed by mobility shift assays. 5 μg of GST-yCTD samples were taken from kinase reactions, then loaded and separated on 8% denaturing Tris-glycine polyacrylamide gels and stained with Coomassie solution. Stained gels were imaged with Gel Doc XR+ Gel Documentation System (Biorad).

### In vitro confocal microscopy

GST-yCTD samples were fluorescently labeled as described in above. Fluorescently labeled GST-yCTD samples were mixed with crowding reagent (dextran) and buffer. Then, 10 μL of samples were directly loaded onto glass slides, covered with 22 mm coverslips. Fluorescent images were acquired with a Nikon W1 Spinning Disk Confocal Microscope with either 60x objective (water immersion) or 100x objective (oil immersion). Fluorescent images were processed using NIS-Elements Viewer (Nikon).

### Differential interference contrast (DIC) microscopy

Wild-type GST-yCTD samples or GST-yCTD variants were mixed with various amounts of dextran, and droplet formation was monitored by DIC microscopy. 10 μL of each sample were applied to slide glass and covered with 22 mm coverslip DIC images were acquired with Nikon eclipse Ni Compound Microscope with 60x objective. DIC images were processed using NIS-Elements Viewer (Nikon).

### In vivo immunofluorescence and microscopy

HeLa and U2OS cells were transfected with MED1-mCherry, Pin1-YFP, SRSF2-mCherry or RPB1aAmr-YFP plasmids for 24 hours before harvest, fixed in 4% paraformaldehyde in PBS for 10 min at room temperature, permeabilization with PBS containing 0.1% Triton X-100 for 10 min at room temperature, then blocked in 2% bovine serum albumin (BSA) in PBS for 1 hours, and incubated sequentially with primary antibodies anti-pSer5 CTD S2 (Millipore, 04-1571, 1:500 dilution), anti-pSer5 CTD S5 (Millipore, 04-1572, 1:500 dilution) or anti-pSer5 CTD T4 (Active Motif, 61361, 1:200 dilution) for overnight at 4°C and Alexa-labeled secondary antibodies (Invitrogen, A11006, A11077, 1:500 and 1:1000 dilution separately) for 1 hours at room temperature with extensive washing. Slides were stained with DAPI (Sigma, MBD0015) and mounted with anti-Fade fluorescence mounting media (Abcam, ab104135). Immunofluorescence images were obtained and analyzed using the Zeiss LSM710 confocal microscope, Nikon spinning disk confocal microscope and ImageJ software.

### Fluorescence recovery after photobleaching (FRAP)

MED1-mCherry were transfected in HeLa cells and Pin1-YFP or SRSF2-mCherry were transfected in U2OS cells for 36 hours and the condensates were photobleached and imaged with a 405_nm laser using Zeiss LSM710 confocal microscope. At each time point, fluorescence intensity within the bleaching spot was divided by the intensity of a neighboring unbleached area of the same size to correct the changes.

### Fluorescence polarization

CTD peptides with double repeats were labeled with fluorescein isothiocyanate (FITC) and purchased from Biomatik. Protein and peptide concentrations were determined according to their absorbance at 280 nm. Fluorescence polarization values were collected on a Tecan F200 plate reader in buffer (50 mM Tris pH 8.0, 300 mM NaCl) at room temperature. Samples were excited with vertically polarized light at 485 nm and at an emission wavelength of 535 nm. Recombinant Pin1, RPRD1B-CID or SCAF4-CID proteins were titrated into a reaction mixture containing buffer supplemented with 100 nM of FITC-peptide. Measurements were taken in triplicates and the experimental binding isotherms were analyzed in GraphPad Prism v8 using a 1:1 binding mode to obtain Kd values.

### Chromatin immunoprecipitation (ChIP) and ChIP-Sequencing

HEK 293T cells were seeded in 15 cm dishes and transfected YFP-RPB1_52xCTD, 26xCTD and mutant plasmids. After 24 h transfection, cells were fixation with 1% formaldehyde for 8 min at room temperature. Crosslinking was quenched with 0.125 M glycine for 5 min. Cells were successively lysed in lysis buffer LB1 (50 mM HEPES-KOH, pH 7.5, 140 mM NaCl, 1 mM EDTA, 10% glycerol, 0.5% NP-40, 0.25% Triton X-100, 1× PI), LB2 (10 mM Tris-HCl, pH 8.0, 200 mM NaCl, 1 mM EDTA, 0.5 mM EGTA, 1× PI) and LB3 (10 mM Tris-HCl, pH 8.0, 100 mM NaCl, 1 mM EDTA, 0.5 mM EGTA, 0.1% Na-deoxycholate, 0.5% N-lauroylsarcosine, 1×PI). Chromatin was sonicated to an average size of ∼200–500 bp using Q800R3 Sonicator (30 s on and 30 s off for 25 min). A total of 5 μg of GFP antibody that was pre-mixed in a 50 uL volume of Dynabeads protein A (Invitrogen) was added to each sonicated chromatin sample with 1% Triton X-100 and incubated overnight at 4°C. The chromatin-bound beads were washed two times with low salt wash buffer (0.1% Na Deoxycholate, 1% Triton X-100, 1 mM EDTA, 50 mM HEPES pH 7.5, 150 mM NaCl), once with high salt wash buffer (0.1% Na Deoxycholate, 1% Triton X-100, 1 mM EDTA, 50 mM HEPES pH 7.5, 500 mM NaCl), once with LiCl wash buffer (250 mM LiCl, 0.5% NP-40, 0.5% Na-Deoxycholate, 1 mM EDTA, 10 mM Tris-Cl pH 8.0) and twice in TE buffer. The chromatin was reverse crosslinked overnight at 65°C with shaking at 750 rpm in cross-linking buffer (1% SDS and 0.1 M NaHCO_3_). After DNA extraction using phenol-chloroform, the DNA was resuspended in 10 mM Tris-HCl pH 8.0. The purified DNA was subjected to qPCR to confirm target region enrichment before moving on to deep sequencing library preparation. For sequencing, the extracted DNA was used to construct the ChIP-Seq library using the NEBNext Ultra II DNA Library Prep Kit, followed by sequencing with an Illumina NovaSeq 6000 system by Novogene.

### RNA isolation, library preparation, and RNA-Sequencing

Total RNA was isolated from HEK 293T cells (at least ∼10^6^ cells/sample) using DirectZol RNA Miniprep kit (Zymo Research, Irvine, CA, product number #R2050). Poly (A) enrichment RNA-Seq was performed by Novogene, mRNA was purified from total RNA using poly-T oligo-attached magnetic beads. After fragmentation, the first strand cDNA was synthesized using random hexamer primers, followed by the second strand cDNA synthesis using dUTP for directional library. The library was checked with Qubit and real-time PCR for quantification and bioanalyzer for size distribution detection. Quantified libraries will be pooled and sequenced on NovaSeq 6000 instrument (paired-end 2×150, 100 cycles). A minimum number of reads was set to 40×10^6^ per sample.

### Analyses of ChIP-Seq data and calculation of Pausing Index (PI)

Initial quality assessment showed high library complexities with low level of duplication. Adapter sequences and low-quality read ends were trimmed off by TrimGalore! v.0.6.7 with default parameters. Paired-end reads were aligned to human reference genome, GRCh38 version, using Bowtie2 v.2.4.5^67^ with default parameters. Mapping stats confirmed high alignment rates (>80-90% of reads aligned concordantly exactly 1 time). Next, coverage bigwig files normalized by Input (IgG-control) were generated out of bam files for every sample using log2 of the number of reads ratio (mapq > 10)^68^. DeepTools v.3.5.1 were also used to prepare score matrices and plotting metagene Pol II ChIP-Seq profiles (over subset of protein-coding genes, n = 19,984 regions). Next, CTD_26x and CTD_52x filtered reads (mapq > 10) were used to call ‘broad’ peaks (p < 0.005) with MACS2 peak caller v.2.2.7.1 keeping one duplicate tag at the exact same location (--keep-dup 1). Obtaining consensus peaksets and occupancy analysis was performed using ‘DiffBind’ v.3.0.15 pipeline^69^ in R. The absolute majority of consensus peaks in both conditions (CTD_26x/52x) reached IDR-threshold (IDR/irreproducible discovery rate < 0.05): 84% and 89% peaks, respectively.

To calculate Pausing Index (PI) as the measure of promoter-proximal pausing of RNA Pol II under CTD_26x vs CTD_52x condition, we used Input-normalized read count files in bigwig format. PI was defined as follows:

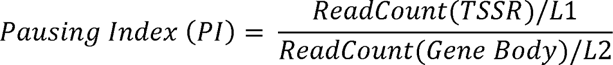

Where TSSR (transcription start site region) is (−50 bp to +300 bp around TSS), and the gene body is (+300 bp downstream of the TSS to +3 kb past the TES). L1 and L2 are the corresponding lengths of the regions^46^. The read densities were calculated using Bwtool (https://github.com/CRG-Barcelona/bwtool, “summary” function) which sums up signal in normalized bigwig files over the defined genomic regions in a bed file containing corresponding to “numerator” and “denominator” coordinates of human protein-coding genes (n = 19,984) derived from gencode hg38 annotation.gtf. Further analysis was conducted in R, where the genes were ranked depending on their average PI across the conditions and then clustered into four groups: G0 cluster with PI = 0, G1 cluster with PI < 25% quartile, G2 cluster with 25% < PI < 75% quartile; and G3 cluster with PI > 75% quartile (most paused genes have PI >=2^46^). ChIP-Seq data was deposited in GEO under the accession number GSE252261.

### Analyses of RNA-Seq data and alternative splicing events (ASE)

As with raw ChIP-Seq reads, adapter sequences and low-quality read ends were trimmed off by TrimGalore! v.0.6.7 with default parameters. Trimmed reads were aligned to human reference genome, GRCh38 version, using HISAT2 fast aligner v.2.2.1 with default parameters, except Reverse (RF) --rna-strandedness. Gencode v38 gtf file was used as annotation gtf. Lastly, mapped fragments were quantified by featureCounts v.2.0.1 in Galaxy^70^.

Differential expression in CTD_26x vs CTD_52x was analyzed using raw unnormalized counts in DESeq2 v.1.30.1 in R; genes with adjusted p-value < 0.05 and |log2FC| > 0.58 were considered as differentially expressed^71^. rMATS turbo v.4.1.2 was employed for detection of alternatively spliced events upon CTD_26x vs. WT-CTD_52x^50^. As input files for rMATS, we used alignment .bam files from HISAT2 mapper (two biological replicates per condition) and gencode v38 annotation gtf. Downstream analysis of rMATS output files containing JCEC counts (Junction Counts and Exon Coverage) was performed in SpliceTools suite^51^. Overrepresentation analysis of gene clusters (gene ontology) was performed using Bioconductor R package ‘clusterProfiler’ v.3.18.1. RNA-Seq data was deposited in GEO under the accession number GSE252260.

### Quantification and Statistical Analyses

Statistical analyses were performed using RStudio v.4.0.5 and GraphPad Prism v9. Two-tailed, independent sample t-test was used to compare the two groups. p < 0.05 was considered significant. For cell survival analysis, oneway ANOVA was performed to determine p-values. Correlations were assessed using two-tailed Pearson r coefficients.

---

**Supplementary Figure 1.**
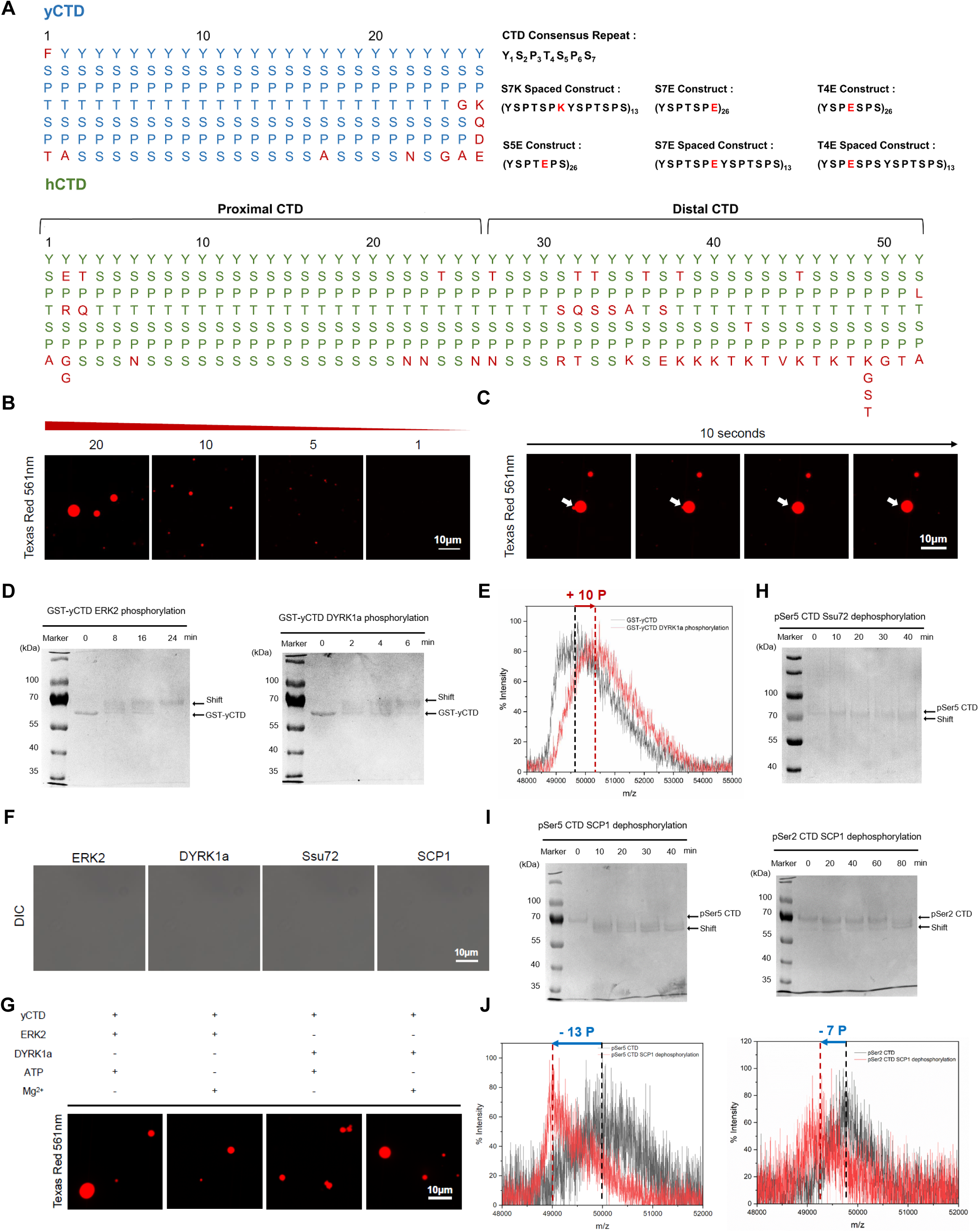
Reversible phosphorylation of the CTD leads to phase transition. (A) The amino acid sequence of RNA polymerase II CTD and its variants used in the study. The CTD sequences of yeast (S. cerevisiae) and human are shown and aligned to display heptad repeats. The sequences of CTD variants are shown right next to the sequence of yeast CTD, and the point of mutations in heptad repeats are highlighted in red. (B) Representative confocal microscopy images of different amounts (μM) of GST-yCTD (Red, Texas Red-X) in the presence of 16% dextran. (C) GST-yCTD droplets (Red, Texas Red-X) show liquid-like fusion upon contact with each other (10 μM GST-yCTD with 16% dextran). (D) SDS-PAGE gel analysis of unphosphorylated CTD samples treated with ERK2 (left) and DYRK1a (right). Change in electrophoretic mobility caused by phosphorylation is shown by gradual shift of protein bands corresponding to GST-yCTD. (E) MALDI-TOF MS spectrum of DYRK1a treated CTD samples from the turbidity assay experiment. (F) Representative DIC image of 10 μM ERK2, DYRK1a, Ssu72, and SCP1 mixed with 16% dextran without GST-yCTD. (G) Confocal microscopy images that represent control experiments of GST-yCTD phosphorylation experiments. In each condition, a component was omitted to control the effect. (H) SDS-PAGE gel analysis of pSer5 CTD samples treated with Ssu72. Change in electrophoretic mobility caused by removal of phosphates is shown by gradual shift of protein bands corresponding to GST-yCTD. (I) SDS-PAGE gel analysis of pSer5 CTD samples (left) and pSer2 CTD samples (right) treated with SCP1. Change in electrophoretic mobility caused by removal of phosphates is shown by gradual shift of protein bands corresponding to GST-yCTD. (J) MALDI-TOF MS spectrum of SCP1 treated pSer5 CTD samples (left) and pSer2 CTD samples (right) from the turbidity assay experiments.

**Supplementary Figure 2.**
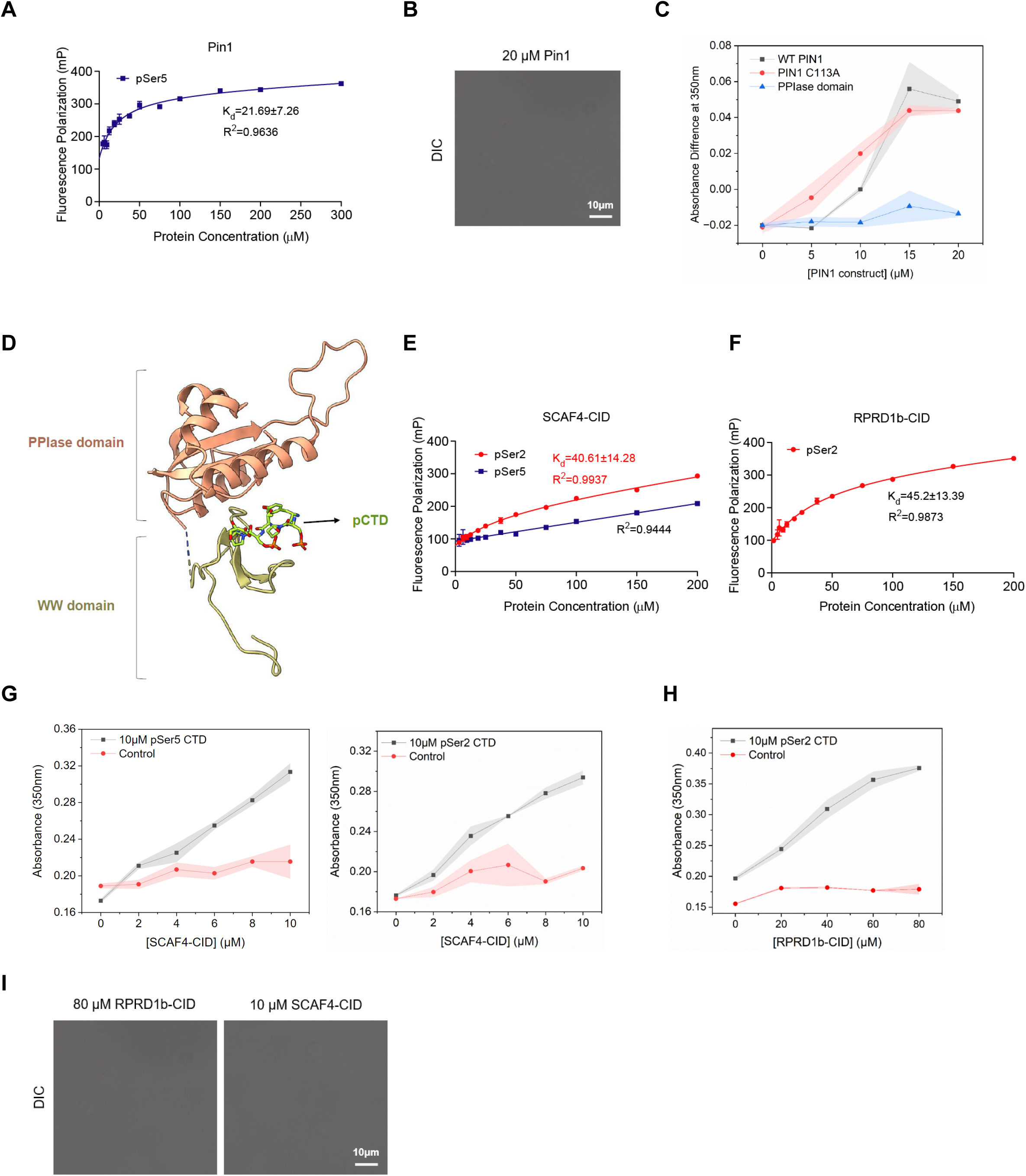
Phospho-specific association of proteins with CTD promotes the reformation of droplets, different CTD condensates remain distinct based on their physical properties. (A) Fluorescence polarization (FP) measurement of the binding of pS5 FITC-labeled CTD peptides containing two repeats to recombinant PIN1. All the experimental isotherms were fitted to 1:1 binding model. Binding assays were performed in triplicate. Error bars indicate standard error of the mean. (B) Representative DIC image of 20 μM PIN1 mixed with 16% dextran without GST-yCTD. (C) Turbidity assay of 10 μM pSer5 CTD mixed with increasing amounts of PIN1 constructs. Each absorbance difference value is obtained by subtracting the blank absorbance obtained from negative control experiments where only PIN1 constructs are mixed with 16% dextran. Data points represent mean values of three replicate experiments, and error bars show the standard error. (D) Crystal structure of PIN1 binding to phosphorylated CTD (Green stick) (PDB code 1F8A). WW domain, represented as yellow-green cartoon, is necessary for substrate recognition of PIN1. PPIase domain (orange cartoon) by itself cannot effectively bind to phosphorylated CTD. (E-F) Fluorescence polarization (FP) measurement of the binding of pS2 or pS5 FITC-labeled CTD peptides containing two repeats to SCAF4 (E) and RPRD1B (F). Experimental isotherms were fitted to a 1:1 binding model. Binding assays were performed in triplicate. Error bars indicate the standard error of the mean. (G-H) Dose-dependent turbidity assay of 10 μM pSer5 CTD and pSer2 CTD treated with different amounts of CTD binding proteins. The sample curve is labeled black while the control is labeled red. Data points represent mean values of three replicate experiments and error bars show the standard error. (I) Representative DIC image of 80 μM RPRD1b, 10 μM SCAF4 mixed with 16% dextran without GST-yCTD.

**Supplementary Figure 3.**
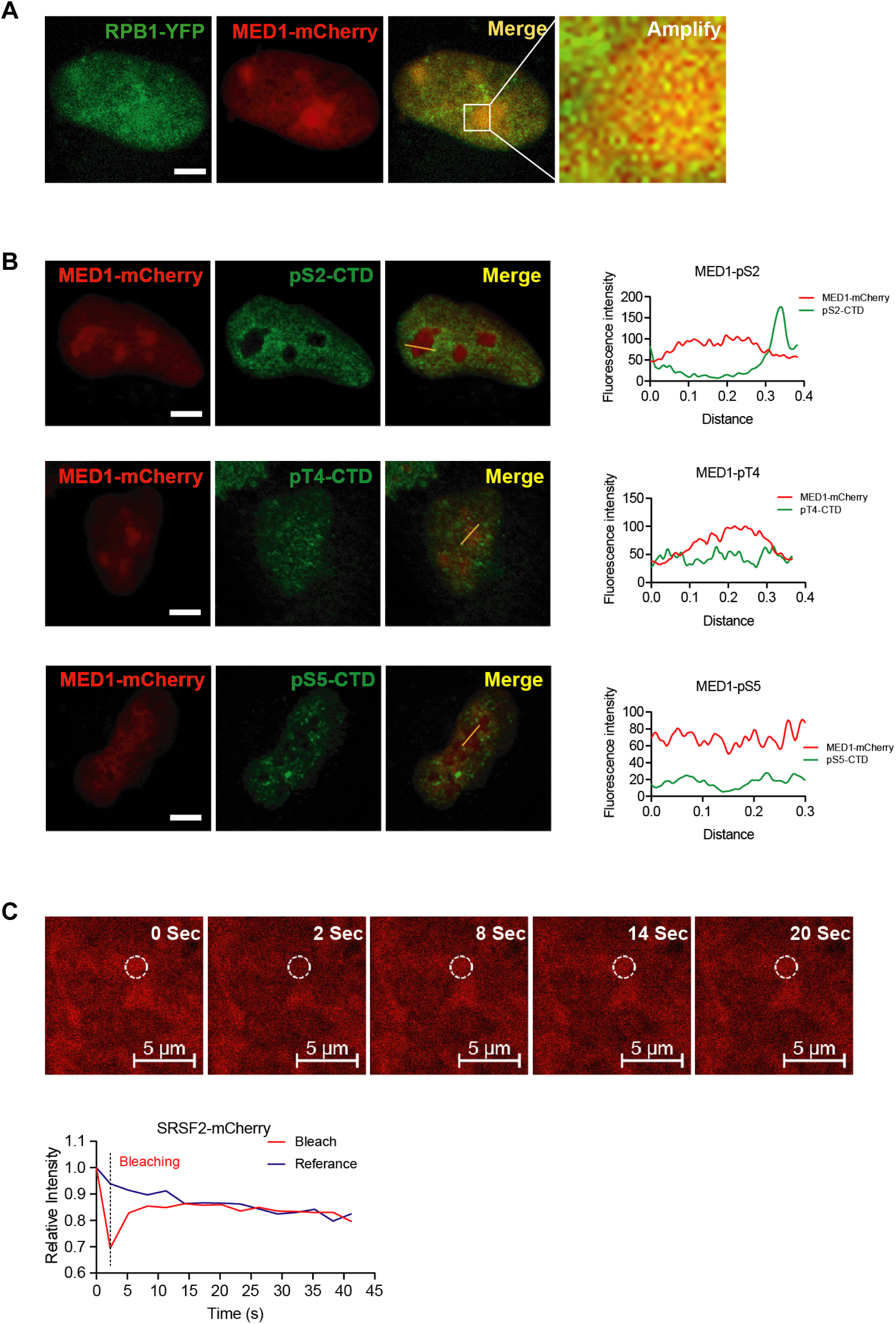
CTD binding proteins colocalized with puncta formed by phosphorylated RNA polymerase II in cells. (A) HeLa cells were transfected with MED1 IDR-mCherry and YFP-RPB1_52xCTD construct and investigated under confocal microscope. Scale bars, 5 μm. (B) HeLa cells were transfected with MED1 IDR-mCherry construct and stained for different phospho-CTD antibodies as indicated. Scale bars, 5 μm. (C) U2OS cells were transfected with SRSF2-mCherry constructs, SRSF2 droplets were detected by FRAP assay in living cells. Scale bars, 5_μm. Changes in the fluorescence intensity of droplets after photobleaching were plotted over time.

**Supplementary Figure 4.**
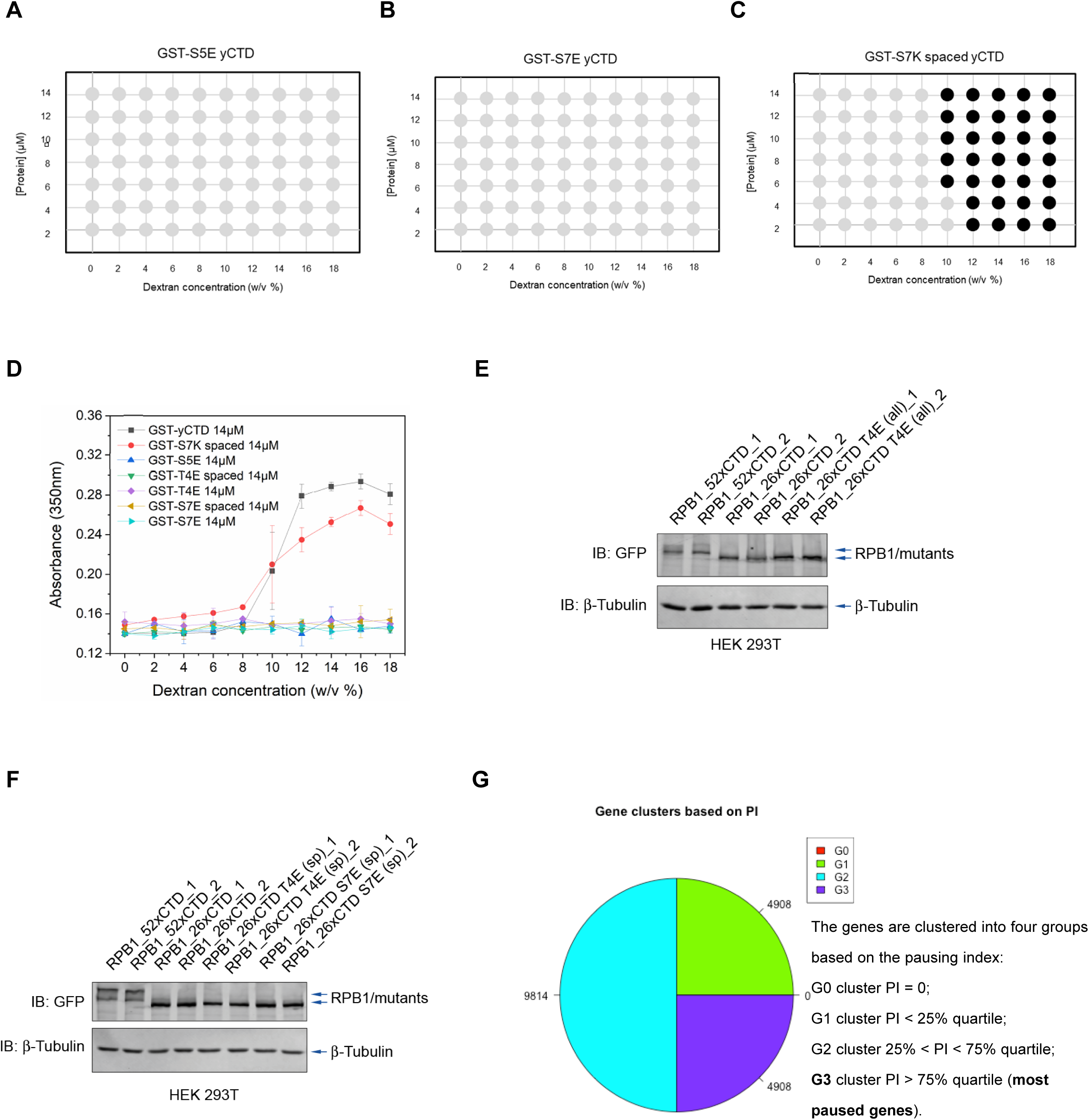
CTD condensation properties in vitro predict genomic locations of RNA polymerase II in vivo. (A) Phase diagram of GST-S5E yCTD. Black filled circles indicate conditions where liquid-liquid phase separation is observed with DIC microscope. (B) Phase diagram of GST-S7E yCTD. (C) Phase diagram of GST-S7K spaced yCTD. (D) Turbidity assay of 14 μM CTD constructs mixed with increasing amounts of dextran. Data points represent mean values of three replicate experiments, and error bars show the standard error. (E-F) Western blotting detected the expression level of YFP-RPB1_52xCTD, 26xCTD and different mutants in Chip-Seq samples. (G) ChIP-Seq signal intensity of genes are clustered into four groups based on the pausing index.

**Supplementary Figure 5.**
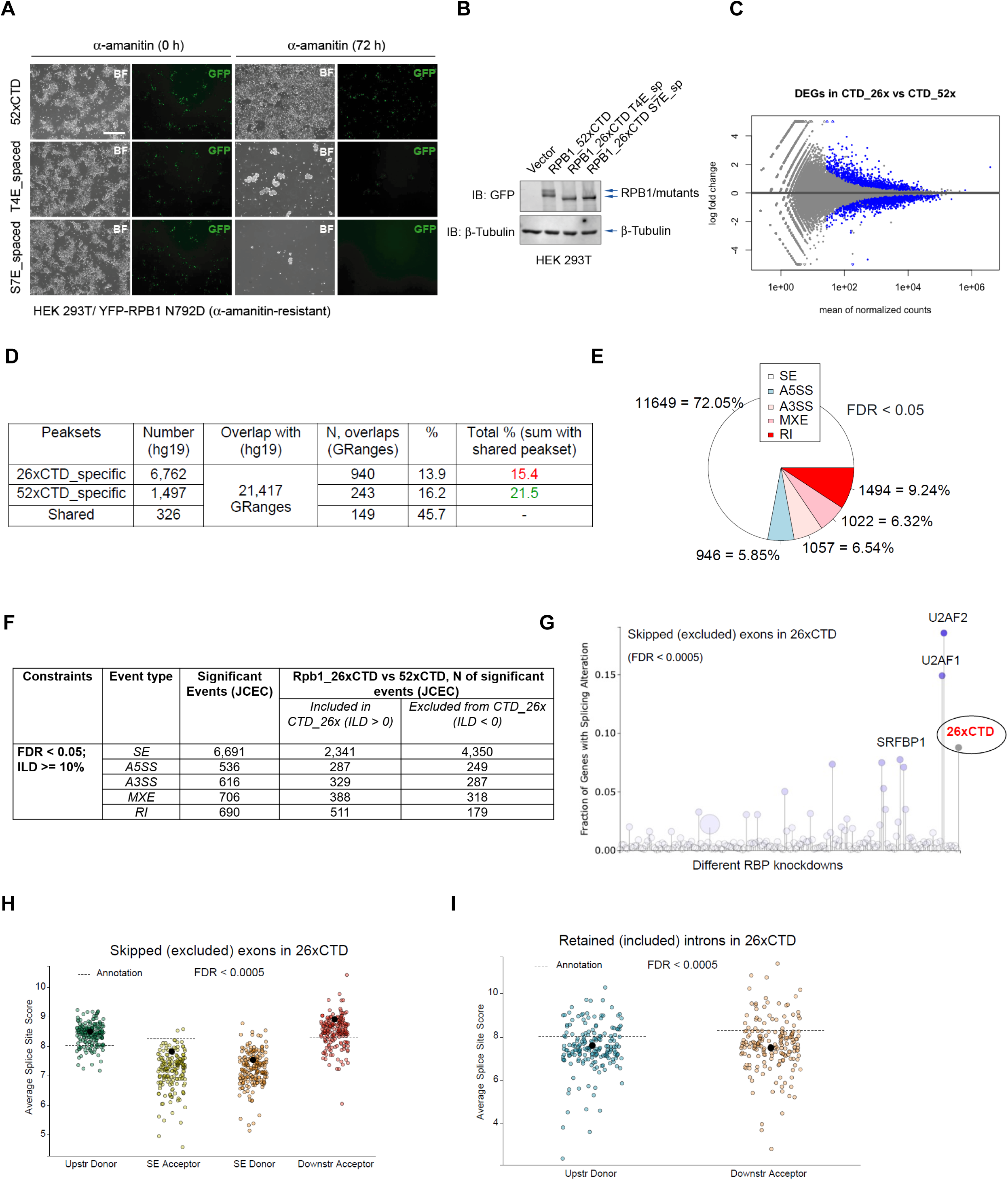
Different CTD variants alter cell survival and alternative splicing. (A) Cell growth of HEK 293T which transfected with RPB1_52xCTD, 26xCTD T4E (spaced) and 26xCTD S7E (spaced) constructs upon α-amanitin administration. (B) Western blotting detected the expression level of RPB1_52xCTD, 26xCTD T4E (spaced) and 26xCTD S7E (spaced) constructs. (C) MA plot showing DEGs upon RPB1_26xCTD compare to RPB1_52xCTD. (D) Calculate overlaps with enhancers of RPB1_26xCTD or RPB1_52xCTD with genes from annotation hg19. (E) Alternative splicing events [in %] detected in RPB1_26xCTD compare to RPB1_52xCTD (with parameters FDR < 0.05). (F) Types and absolute numbers of alternative splicing events (ASE) that were significantly different in RPB1_26xCTD vs. RPB1_52xCTD cells (with parameters FDR < 0.05; ILD, inclusion level difference, ≥ 10%). (G) Transcriptome footprint of skipped exons (SE) in RPB1_26xCTD overexpression, splicing factors and RBP knockdowns. (H-I) Average splice site scores for skipping exons (H) and retaining introns (I) in RPB1_26xCTD. Dashed lines provide reference scores for potential skipping and retaining configurations derived from an input annotation file.

